# Detyrosination enrichment on microtubule subsets is established by the interplay between a stochastically-acting enzyme and microtubule stability

**DOI:** 10.1101/2022.09.29.510213

**Authors:** Qing Tang, Sebastian Sensale, Charles Bond, Andy Qiao, Siewert Hugelier, Arian Arab, Gaurav Arya, Melike Lakadamyali

## Abstract

Microtubules in cells consist of functionally diverse subpopulations carrying distinct post-translational modifications (PTMs). Akin to the histone code, the tubulin code regulates a myriad of microtubule functions ranging from intracellular transport to chromosome segregation. Yet, how individual PTMs only occur on subsets of microtubules to contribute to microtubule specialization is not well understood. In particular, microtubule detyrosination, which is the removal of the C-terminal tyrosine on α-tubulin subunits, marks the stable population of microtubules and modifies how microtubules interact with other microtubule-associated proteins to regulate a wide range of cellular processes. Previously, we found that, in certain cell types, only a small subpopulation of microtubules is highly enriched with the detyrosination mark (∼30%) and that detyrosination spans most of the length of a microtubule, often adjacent to a completely tyrosinated microtubule. How the activity of a cytosolic detyrosinase, Vasohibin (VASH) leads to only a small subpopulation of highly detyrosinated microtubules is unclear. Here, using quantitative super-resolution microscopy, we visualized nascent microtubule detyrosination events in cells consisting of 1-3 detyrosinated α-tubulin subunits after Nocodazole washout. Microtubule detyrosination accumulates slowly and in a disperse pattern across the microtubule length. By visualizing single molecules of VASH in live cells, we found that VASH engages with microtubules stochastically on a short time scale suggesting limited removal of tyrosine per interaction, consistent with the super-resolution results. Combining these quantitative imaging results with simulations incorporating parameters from our experiments, we propose a stochastic model for cells to establish a subset of detyrosinated microtubules via a detyrosination-stabilization feedback mechanism.

## Introduction

Microtubules experience numerous posttranslational modifications (PTMs), including polyglutamylation, acetylation and detyrosination, which segregate them into distinct subpopulations with diverse mechanical and functional properties [1, 2]. Little is known about how certain microtubule PTMs are present or enriched on only subsets of the total microtubule population. Microtubules are dynamic cylindrical polymers assembled from α, β-tubulin heterodimers, the C-terminal tails of which are exposed to the cytosol and are particularly enriched with PTMs. These cytosolic tubulin tail modifications allow direct tuning of interactions between microtubules and many microtubule-associated proteins (MAPs). Most α-tubulin isoforms encode a C-terminal tyrosine, which can be removed by the recently discovered carboxypeptidase Vasohibin (VASH) [3, 4] and another newly discovered tyrosine carboxylpeptidase MATCAP [5]. It has been shown that both in the mouse brain and in vitro cell lines, VASH is the major detyrosinase that accounts for the vast majority of detyrosination (∼ 75% in mouse brain [3]), whereas MATCAP only contributes a modest amount of detyrosination [4, 5]. Detyrosination changes how certain MAPs interact with the C-terminal tail of α-tubulins on microtubules. For example, MAPs containing the cytoskeleton-associated protein glycine-rich (CAP-Gly) domain directly recognize the tyrosine residue of the EEY/F motif at the C terminus of α-tubulin and preferentially bind to tyrosinated tubulin [6–11]. Binding of certain motor proteins is also regulated by the detyrosination state of the microtubule [7, 12–17]. Detyrosination is a particularly important microtubule PTM as numerous cellular processes are regulated by microtubule detyrosination, including mitosis/meiosis, intracellular transport, autophagy, neurogenesis, and cardiomyocyte contractility [2, 5, 15–24]. Detyrosinated microtubules are known to persist much longer than tyrosinated microtubules in live cells [25–27]. It has long been proposed that this stability is not due to detyrosination itself altering the intrinsic dynamics of microtubules, but the change in interaction with many MAPs that collectively alter the microtubule dynamics [28–30]. Indeed, a recent in vitro study employing recombinantly expressed α-tubulin lacking the C-terminal tyrosine demonstrated that these detyrosinated microtubules are not intrinsically more stable than tyrosinated microtubules without the influence of MAPs [8].

Previously, using super-resolution microcopy, we classified microtubule sub-populations based on their PTMs in epithelial cell types and found that detyrosination only occurs on ∼30% of total microtubules, while acetylation is much more predominant and constitutes ∼70% of total microtubules [16]. We further showed that lysosomes and autophagosomes are enriched on this small subset of detyrosinated microtubules and that this enrichment is important in promoting their fusion. Hence, building a small subpopulation of detyrosinated microtubules that can spatially regulate organelle positioning seems to be crucial for cellular processes like autophagy. It has also been shown that in neuronal dendrites, tyrosinated microtubules are concentrated on the outer portion of the dendrite, spatially segregated from acetylated (and also presumably detyrosinated) microtubules [31], and the two populations were proposed to mediate motor transport in opposite directions [15]. These results overall show that in several cell types including epithelial cells and neurons, detyrosinated microtubules constitute a distinct subset of all microtubules to regulate diverse cellular processes.

Because VASH prefers polymerized tubulin as a substrate compared to free tubulin dimers [3, 4, 32, 33], detyrosination occurs more readily once the microtubules are assembled. On the other hand, the tubulin tyrosine ligase (TTL) adds back the tyrosine primarily on detyrosinated free tubulin dimers [34], therefore detyrosination is likely stably present on assembled microtubules and α-tubulin does not revert to tyrosinated form until the tubulin subunit is released back to the cytosolic pool upon microtubule depolymerization. It is thought that TTL stays associated with the tubulin pool in the cytosol and sequesters tubulin dimers [34], consistent with early work suggesting that the cytosolic tubulin pools are largely tyrosinated [35] and that nascently rescued stretches of microtubules are also tyrosinated [36]. Additionally, early work showed that when detyrosinated tubulin dimers were injected into the cells in which most of microtubules were depolymerized by Nocodazole, they became tyrosinated relatively quickly [37]. Consequently, overall detyrosination levels respond to the change of the balance between free tubulin dimers and microtubules, i.e., total detyrosination decreases markedly upon microtubule disassembly, and increases upon microtubule stabilization [32, 33, 35]. However, the long-standing question remains as to how a subset of highly detyrosinated microtubules emerges adjacent to completely tyrosinated ones through the action of a cytosolic detyrosinase that predominantly acts on polymerized microtubules. Multiple models are possible, including cooperativity among the VASH enzymes especially when enzyme concentration is limited, processive motion or facilitated diffusion of the VASH enzyme along the microtubule lattice, or preferential accumulation/recruitment of the VASH enzyme to a sub-set of microtubules.

To address this long-standing question, we investigated how cells establish and propagate detyrosination on a subset of microtubules using a combination of multicolor DNA Point Accumulation in Nanoscale Topography (DNA-PAINT) [38] super-resolution microscopy, single molecule tracking and computational modeling. Our experimental results coupled with stochastic modeling using parameters extracted from our experiments suggest a mechanism where selective enrichment of detyrosination is a function of enzyme concentration and arises from the positive feedback between detyrosination and microtubule stabilization.

## Results

### Nascent detyrosination appears as disperse and isolated puncta along microtubules and increases slowly over time

To understand how detyrosination progressively builds up in cells, we treated BSC-1 epithelial cells with Nocodazole for three hours, which leads to almost complete microtubule disassembly [36], and washed out the compound to allow the microtubules to reassemble over time. Cells were fixed every 30 minutes post washout, stained with antibodies against β-tubulin and detyrosinated α-tubulin and imaged using multi-color DNA-PAINT super-resolution microscopy [38]. Similar to our previous study, in untreated cells, we observed detyrosination enriched on a subset of microtubules while many adjacent microtubules were completely tyrosinated (**Figure 1**). Upon Nocodazole treatment, on the other hand, the majority of microtubules, including detyrosinated microtubules disappeared as expected (**Figure 1**) [32, 35–37], which allowed us to monitor how nascent detyrosination re-emerges.

**Figure 1:**
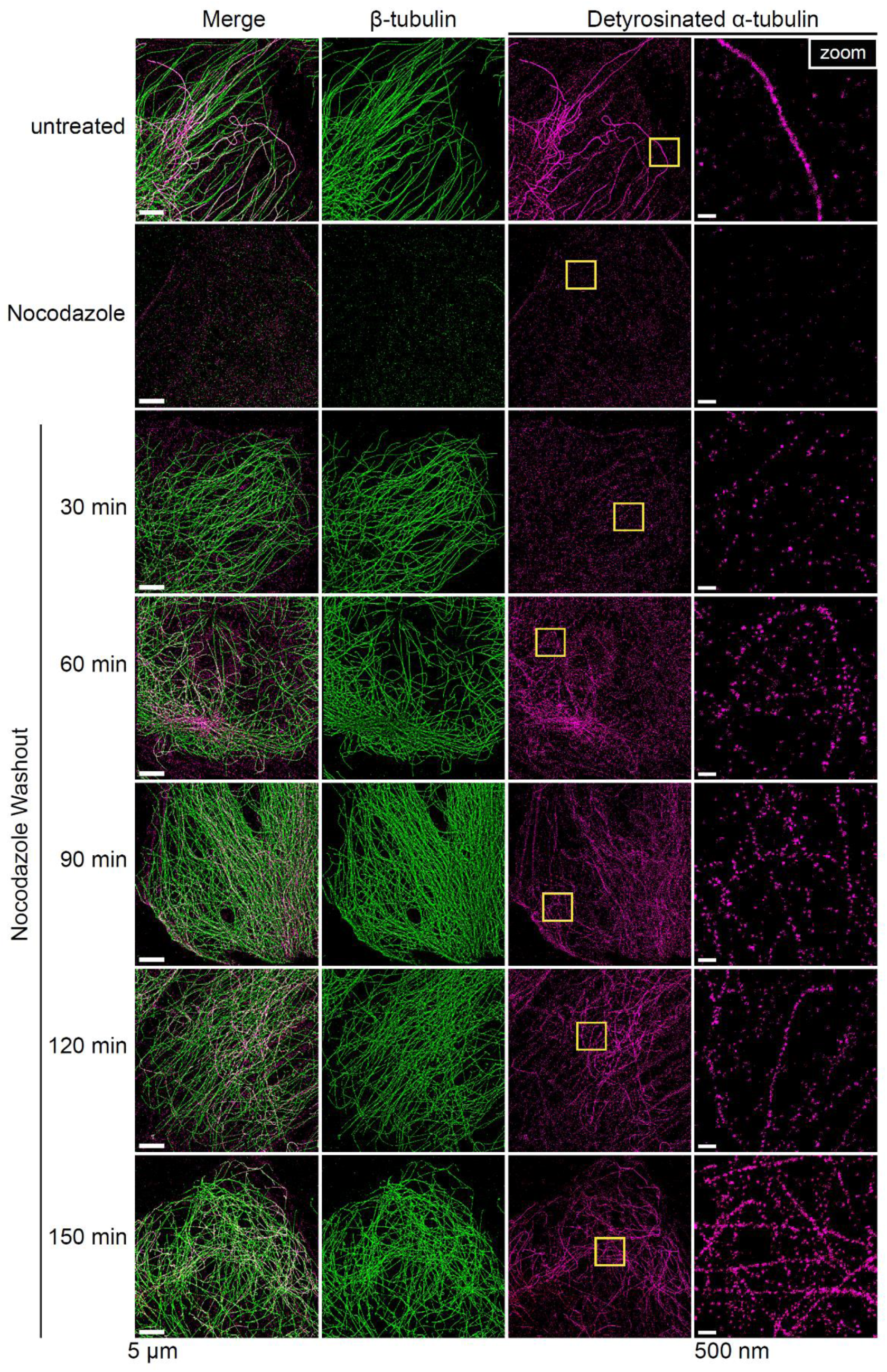
Super-resolution reveals detyrosination is established gradually from small, disperse puncta over time. Representative two-color DNA-PAINT images showing detyrosinated α-tubulin (magenta) and β-tubulin (green) in fixed BSC-1 cells that are treated with 0.1% DMSO (untreated), or 33 µM Nocodazole for 3 hr at 37 °C, and at 30-, 60-, 90-, 120-, 150-min after the Nocodazole washout. Yellow boxes indicate the regions selected for the zoom images on the right most column. Scale bars, 5 µm. Scale bars in zoom, 500 nm.

At 30 min post washout the majority of microtubules were already re-assembled (**Figure 1**), whereas detyrosination was largely absent on these re-assembled microtubules. However, close examination of the super-resolution images showed that nascent detyrosination events were detectable as sparse, small puncta on some regions of microtubules (176 ± 98 nm nearest neighbor distance between puncta borders), with a median area of 1326 ± 290 nm^2^ and a median length of 85 ± 12 nm (**Figures 2A-C**). At 60-90 min after washout, the frequency of the discrete detyrosinated puncta increased. Accordingly, the median distance between them decreased (163 ± 40 nm at 60 min and 109 ± 19 nm at 90 min) (**Figure 2C**), with only a slight increase of the median detyrosination length (86 ± 9.5 nm at 60 min and 98 ± 8.5 nm at 90 min) (**Figure 2B**), and the median area (1476 ± 329 nm^2^ at 60 min and 1933 ± 237 nm^2^ at 90 min) (**Figure 2A**). These results suggest that the puncta are isolated, stochastic nascent detyrosination events rather than growth of established detyrosination sites. At later time points (120-150 min), we found that the distance between detyrosinated puncta did not change dramatically (**Figure 2C**) while detyrosination median length (114 ± 15 -130 ± 22 nm) (**Figure 2B**), and median area (2454 ± 769 - 3174 ± 972 nm^2^) (**Figure 2A**) continued to increase, which is consistent with the puncta merging together as their frequency increases to generate larger stretches of detyrosinated microtubules (**Figures 1 and 2A-C**).

**Figure 2:**
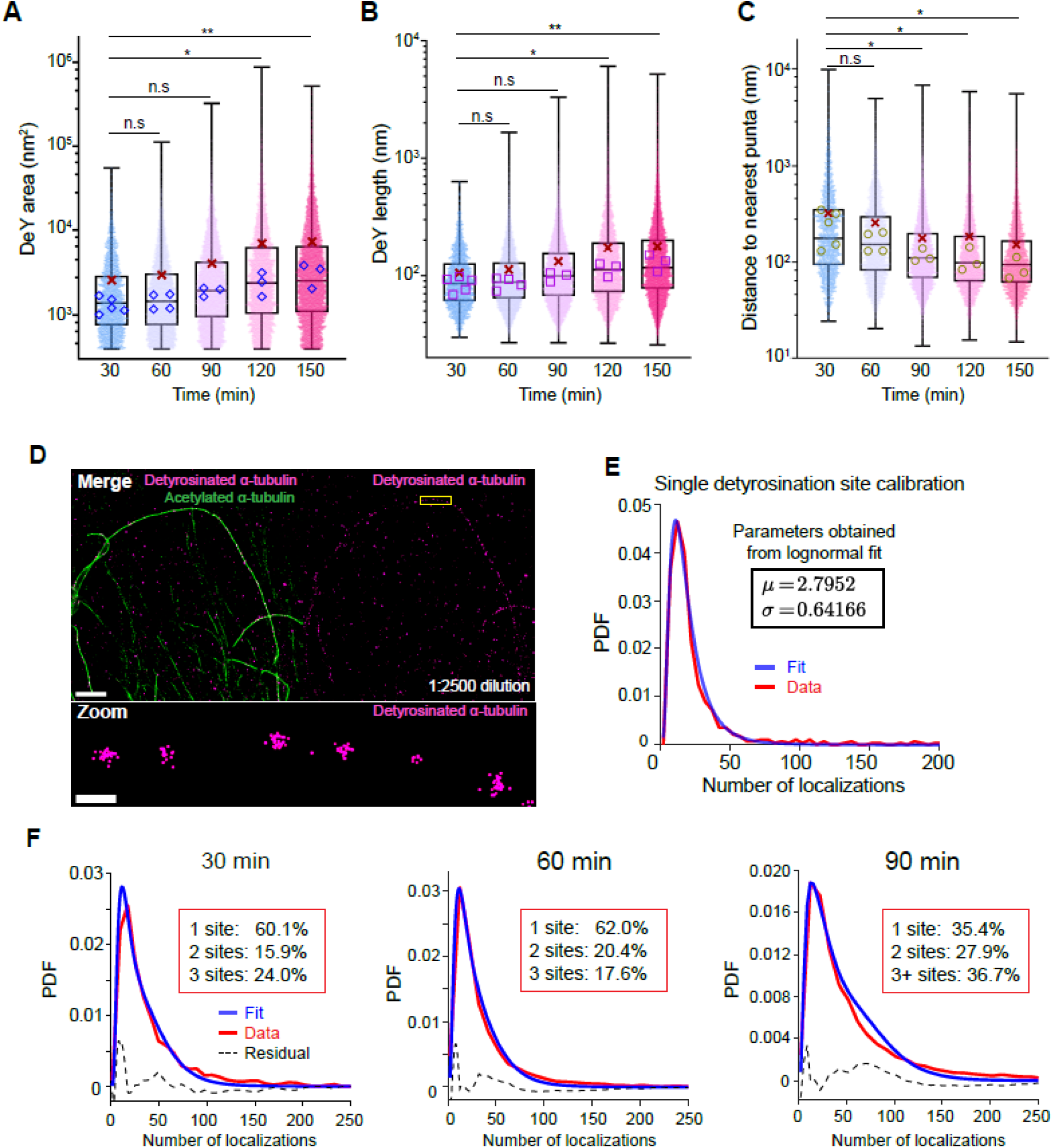
Diffraction limited, nascent detyrosinated puncta correspond to 1-3 detyrosination sites accumulate slowly over time and merge into larger detyrosinated areas. Superplots showing quantitative measurement of detyrosination cluster area (**A**), length (**B**), and the distance to the nearest neighbor cluster (**C**) at each time point after Nocodazole washout shown in Figure1. Many detyrosination events have the appearance of small puncta at early time points (30-min to 60-min), but appear larger and closer to each other at later time points. The measurement of individual puncta at each time point were pooled from 3-5 independent experiments. The scatter plot was overlaid with box and whisker showing 25^th^, 75^th^ percentiles as well as the median values corresponding to each independent experiment (open blue diamonds in (**A**), open purple square in (**B**) and open brown circle in (**C**)). The red crosses indicate the mean values of all data points from each time point. For the statistics of each time point after Nocodazole washout 30 min, *n* = 2204 puncta, *N*=15 cells from 5 experiments; 60 min, *n* = 7846 puncta, *N* =17 cells from 4 experiments; 90 min, *n* = 10686 puncta, *N* = 17 cells from 3 experiments; 120 min, *n* =5957 puncta, *N* = 13 cells from 3 experiments; 150 min, *n* = 7631 puncta, *N* = 9 cells from 3 experiments. One-way ANOVA and Dunnett’s multiple comparison were performed on median values from individual experiments. n.s, not significant, \*\**P* < 0.01, \**P* < 0.05, *P* > 0.05. (**D**) A representative two-color DNA-PAINT super-resolution image used for the purpose of single-detyrosination site calibration, showing detyrosinated α-tubulin and acetylated α-tubulin in untreated BSC-1 cells. Detyrosination appears as small puncta instead of being continuous because the primary antibody was used at 1 to 2500 dilution, a concentration that is 25-fold lower than the saturating amount used in Figure 1. Scale bar in the merge, 2 µm, scale bar in zoom, 200 nm. (**E**) The number of localizations per puncta in images from (**A**) was plotted as a histogram (red) and fitted to a lognormal calibration function (blue, see Methods). The mean (µ) and variance (σ) extracted from the fit are shown. A total of 559 puncta from two experiments were pooled. (**F**) Percentage of puncta having single-, double-, triple or more detyrosinated sites was obtained by the fit of the experimental data for the number of localizations per puncta (red) to a weighted convolution of single, double, triple (or more) calibration functions. The weights were extracted as the percentages. Black dotted lines show the residuals from the fit.

We also observed that the anti-detyrosinated α-tubulin antibody stained non-specifically and produced diffuse background puncta outside of microtubules. During our analysis, we excluded puncta that have a density beyond two-standard deviation from the average as these very dense puncta tend to correspond to imaging artefacts. To determine the contribution from the non-specific background signal to what we considered to be the nascent detyrosinated puncta overlapping with microtubules, we measured the false co-localization percentage of detyrosinated puncta with microtubules by rotating the two images with respect to one another (**Figure S1** and Methods). This analysis showed that the false co-localization percentage was on average only 9-20% compared to the real co-localization (**Figure S1** and Methods), with the higher false co-localization percentage corresponding to the early time points (30-90 min), when detyrosinated puncta were most sparsely located on microtubules. Given that the false co-localization percentage is overall low, especially between 120-150 minutes, we do not expect the non-specific background signal to influence our results.

### Detyrosinated puncta at early time points constitute 1-3 detyrosinated α-tubulin subunits

We next wondered whether the isolated small puncta represent single or multiple detyrosinated α-tubulin subunits, which is a critical question to address for a mechanistic understanding of detyrosination propagation. Since the spatial resolution of our DNA-PAINT images is ∼20-30 nm, whereas the size of a tubulin hetero-dimer is ∼8 nm, we cannot directly resolve multiple detyrosination events happening on adjacent α-tubulin subunits. In single molecule localization microscopy (SMLM), including DNA-PAINT, images of a single protein (or fluorophore) resemble a small cluster of localizations with a broad distribution due to repeated “blinking” and non-stoichiometric antibody labeling [39, 40]. We previously showed that, if the distribution of the number of localizations corresponding to a single protein target is known, it can be fit to a lognormal distribution to obtain a calibration function [39–41]. The calibration function can then be used to deduce the number of protein targets in the actual super-resolution clusters or puncta. To obtain a calibration function for a single detyrosination site, we carried out an antibody titration experiment (1:1000, 1:2500, 1:5000 dilution) to determine a primary antibody concentration that is low enough to sparsely detect detyrosination signal from individual α-tubulin subunits in untreated BSC-1 cells (**Figures 2D** **and S2**). This antibody titration approach has previously been successfully used to estimate protein copy number in super-resolution images by us and others [39–42].

In contrast to images stained with saturating amounts of primary antibodies (1:100) that show a continuous detyrosination signal along the length of the microtubule (**Figure 1, top row**), detyrosination under dilute antibody labeling conditions appeared as sparse, small puncta (**Figure 2D** **and Figure S2**). At the lowest concentration used (1:5000 dilution), there was no detectable detyrosination signal on microtubules (**Figure S2A**). We could detect detyrosination signal with 1:2500 and 1:1000 dilution (**Figures S2B and S2C**), with the 1:1000 dilution generating detyrosinated puncta that contained a slightly higher number of localizations (median 23 vs 16) that were more closely spaced together (median distance 147 nm vs 386 nm) than the 1:2500 dilution (**Figure S2D**). We thus used the sparser 1:2500 dilution as the calibration data for a single detyrosination site. We plotted the frequency distribution of the number of localizations in each detyrosinated puncta obtained from the calibration images and fit it to a lognormal distribution to obtain the mean (µ) and variance (σ) (**Figure 2E**). We then fit the frequency distribution of the number of localizations from the Nocodazole wash out experiments to a linear convolution of the lognormal calibration function to estimate the number of detyrosinated subunits in these images (**Figure 2F**). Our results showed that the puncta at early time points after Nocodazole washout corresponded to 1-3 detyrosinated sites and the proportion of single, double, and triple detyrosinated sites changes over time. In particular, the puncta at 30-60 min after Nocodazole washout were predominantly (∼60%) single detyrosinated sites with only a minor population of 2 and 3 detyrosinated sites (**Figure 2F**). At 90 min, the proportion of puncta containing single detyrosinated sites decreased and the proportion of puncta containing 3 or more detyrosinated sites increased (**Figure 2F**). These results further suggest that nascent detyrosination occurs at a limited (mostly single) number of subunits sparsely and sporadically distributed along the length of the microtubule.

Previously, we showed that the majority of detyrosinated microtubules are also acetylated, although acetylated microtubules constitute a much higher proportion of the total microtubule network [16]. Acetylation occurs on Lys40 on α-tubulin, which is located in the microtubule lumen. To compare the kinetics of acetylation to those of detyrosination, we repeated the Nocodazole washout experiments and stained for acetylated tubulin. The recovery of acetylation after Nocodazole washout occurred much faster than detyrosination. We observed a significant amount of long stretches of acetylation along microtubules at 15 min post Nocodazole washout, and by 30-60 min, a large proportion of microtubules were already acetylated along their entire length (**Figure S3**). These results are consistent with previous reports [43] and in contrast to our results with detyrosination, which starts at sparse, limited sites and accumulates with much slower kinetics in cells.

### Single molecules of VASH bind to microtubules stochastically and show distinct mobility modes

Our super-resolution data suggest that detyrosination occurs stochastically and on only a few subunits at a time on microtubules. To further support this mechanism, we turned to single molecule imaging of the major carboxypeptidase, Vasohibin (VASH) [3, 4], in live cells. Mammals have two isoforms of VASH, VASH1 and VASH2, both of which constitutively complex with the small Vasohibin-binding protein (SVBP) [44] which is a 66-amino acid peptide containing mainly an alpha helix. The two VASH isoforms share conserved structures, including their catalytic cores, albeit recent Cryo-EM results showed some differences in their microtubule binding pose [32, 45].

To determine how individual VASH proteins interact with microtubules in live cells, we co-expressed human SVBP-Myc-FLAG and human VASH1 or VASH2 with a C-terminal Halo Tag from a lentiviral vector in the human U2OS cells, the microtubules of which are largely tyrosinated [46]. Detyrosination significantly increased upon ectopic WT VASH expression compared to uninfected cells (**Figure 3A**), demonstrating that the VASH-Halo/SVBP complex is active. To track single molecules of VASH, we labeled Halo-tag with its ligand conjugated with JF549 [47]. We further labeled microtubules in live U2OS cells with Tubulin tracker (**Figure S4A**) and used the microtubule images as a mask to discard cytosolic VASH molecules that did not overlap with microtubules and only analyzed those trajectories that substantially overlapped with labeled microtubules (**Figures 3B** **and S4B**). As the microtubule image was recorded as a snapshot prior to the single molecule tracking and the end of the single molecule movie is shifted in time by 130 seconds from the microtubule snapshot, we checked for the stability of the microtubules during this time period (**Figure S4A**). Most regions of the microtubule network overlapped but some microtubules showed a modest amount of shift in this time period. Hence, to account for the shift, we considered trajectories that had a minimum of 20% overlap with the microtubule mask to be microtubule associated (Methods).

**Figure 3:**
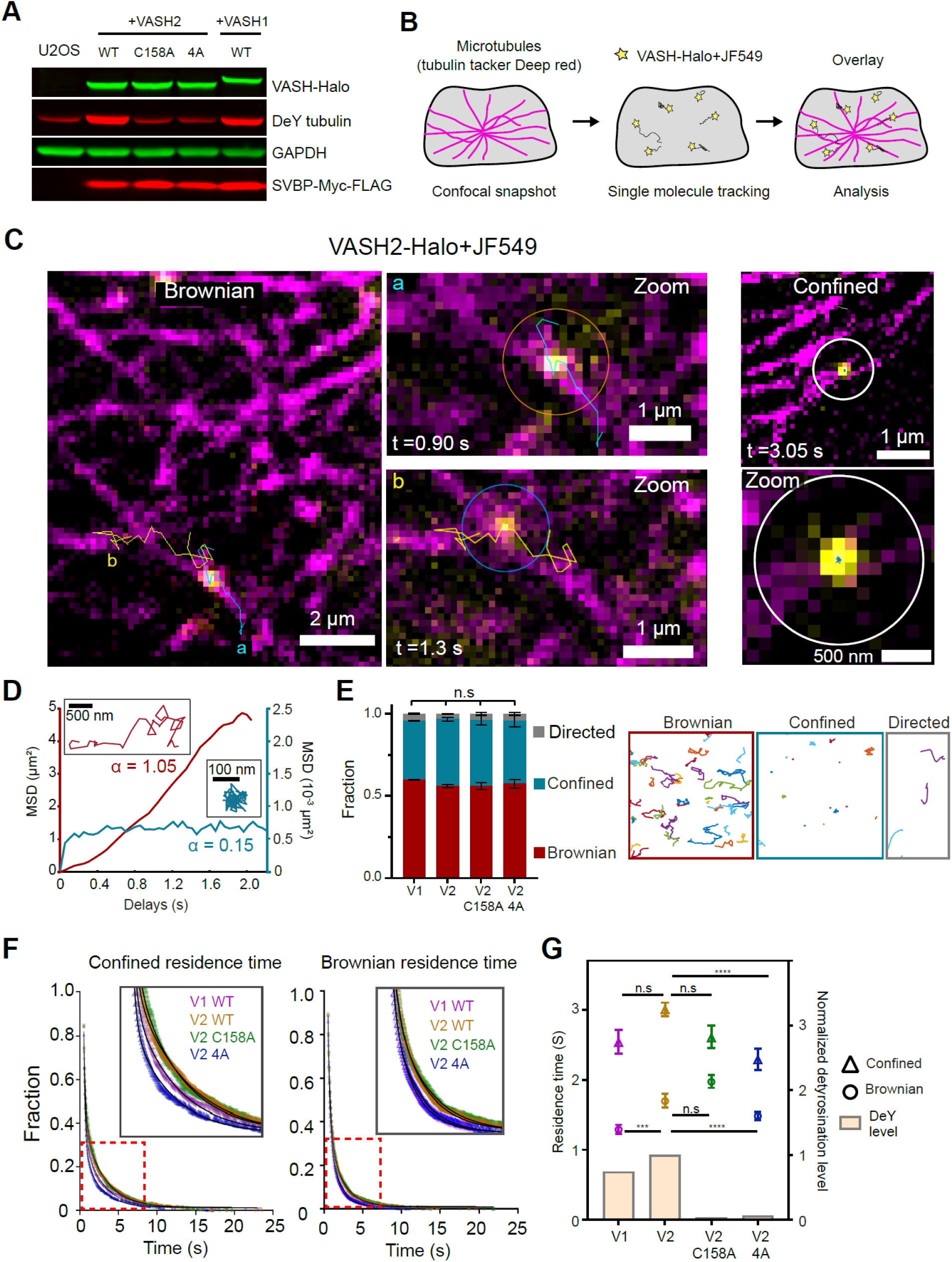
Single molecule tracking reveals VASH briefly interacts with microtubules and catalyzes 1-2 detyrosination events at a time. (**A**) A Western blot showing the ectopic expression of VASH1 WT-, VASH2 WT-, V2 C158A-, and V2 4A-Halo in U2OS cells and the corresponding detyrosination levels (anti detyrosinated α-tubulin). GAPDH was used as a housekeeping protein as a loading control. The detyrosination level significantly increased upon VASH1-Halo WT or VASH2-Halo WT expression. (**B**) Schematic showing the experimental and analysis workflow. (**C**) Representative trajectories showing Brownian and confined motions, each from a single JF549 labeled VASH2-Halo molecule. The trajectories overlapped with the confocal images of microtubules labeled with Tubulin Tracker Deep red (magenta) in live U2OS cells. The duration(t) of each trajectory is indicated. (**D**) Example MSD curves showing a diffusive motion (red) and a confined motion respectively. Insets: the corresponding trajectories. The y-axes are color-coded as the trajectories and scaled to the MSD range of each motion type. The α exponent for each curve is shown. (**E**) Motion classifications for VASH1, VASH2 and VASH2 mutants. The fractions of confined, Brownian and directed motions are plotted as mean ± range (total of 4047, 2992, 2858, and 3974 trajectories pooled from 38 cells, 37 cells, 39 cells and 37 cells for VASH1, VASH2, V2 C158A, and V2 4A respectively, N= 2 experiments). Representative trajectories for each motion type are shown on the right (**F**) Double exponential fits for extracting residence time for confined or Brownian motions for each VASH variant. The insets show zoomed regions (red boxes) of the fits. (**G**) Residence time for confined and Brownian motions extracted from the long time constant of the exponential fit and corrected for photobleaching. Error bars correspond to the 95% confidence interval. The detyrosination level was calculated as a normalized value against the detyrosination level of VASH2 WT, using the western blot in (**A**). Krauskal-Willis test with Dunn’s multiple comparison test was performed on all trajectories from each condition. *** *P*<0.001, **** *P*<0.0001, *P> 0.05*, ns, not significant.

As previously observed, VASH was predominantly cytosolic (**Figure S5**) [3, 4]. However, those trajectories that overlapped with the microtubule mask showed two types of behavior. A proportion of VASH molecules exhibited diffusive behavior on microtubules, sometimes moving bi-directionally for short time periods along a single microtubule (**Figures 3C** **and S4C, Supplementary Movies S1 and S2**). In some cases, we observed prolonged diffusion of VASH along microtubules, in particular in regions where multiple microtubules crisscrossed (**Figure S4D and Supplementary Movies S3 and S4**). A second population of VASH molecules exhibited highly confined motion at a given spot on microtubules for a short period of time before diffusing out of focus (**Figures 3C** **and S4C, Supplementary Movies S5 and S6**). In some select cases, we observed switching behavior between Brownian and confined motion in a single trajectory (**Figure S4E, Supplementary Movie S7**). We carried out mean square displacement analysis for trajectories that contained at least 10 frames (or 0.5 s in tracking time) and classified the motions of VASH WT enzyme molecules into three categories: confined (having an MSD alpha coefficient ≤ 0.8), Brownian (having an MSD alpha coefficient between 0.8-1.5) and directed (having an MSD alpha coefficient ≥ 1.5). The directed trajectories constituted a minimal proportion of all trajectories (2-3%) (**Figures 3D and 3E, Table 1**). These trajectories were also mostly short lived and are likely due to tracking errors or mis-classification due to limitations of the MSD analysis. Interestingly, roughly equal proportion of VASH molecules exhibited Brownian (59.7%, 56.1% for VASH1 and VASH2 respectively) or confined motion (35.9%, 40.6% for VASH1 and VASH2 respectively) on microtubules (**Figure 3E**, **Table 1**).

**Table 1.** Residence time and percent population of motion classes.

| VASH WT and mutants | Brownian |  |  |  | Confined |  |  |  |
| --- | --- | --- | --- | --- | --- | --- | --- | --- |
|  | Percent of total population | Residence time (s) | 95% confidence interval (s) |  | Percent of total population | Residence time (s) | 95% confidence interval (s) |  |
|  |  |  | upper limit | lower limit |  |  | upper limit | lower limit |
| VASH 1 WT | 59.7 | 1.289 | 1.357 | 1.227 | 35.9 | 2.533 | 2.708 | 2.375 |
| VASH2 WT | 56.1 | 1.695 | 1.797 | 1.604 | 40.6 | 3.002 | 3.101 | 2.910 |
| VASH2 C158A | 56.2 | 1.971 | 2.065 | 1.885 | 40.0 | 2.601 | 2.774 | 2.450 |
| VASH2 4A | 58.0 | 1.482 | 1.544 | 1.425 | 37.5 | 2.286 | 2.446 | 2.143 |
| VASH1 WT (washout) | 51.3 | 1.336 | 1.393 | 1.284 | 45.2 | 1.376 | 1.458 | 1.302 |
| VASH2 WT (washout) | 53.7 | 1.703 | 1.870 | 1.558 | 42.5 | 2.227 | 2.333 | 2.128 |

All residence times shown here are corrected against photobleaching (see Methods and Figure S4).

To determine the dwell time of the VASH molecules on microtubules, we fit the residence time for Brownian and confined trajectories to a double exponential. We took the long time constant after photobleaching correction (**Figure S4F** and see Methods) to represent the specific interaction of VASH with microtubules as is standard in single molecule tracking experiments [48–50]. The Brownian residence time was 1.29 s for VASH1, and 1.70 s for VASH2, and the confined residence time was 2.53 s and 3.00 s for VASH1 and VASH2, respectively (**Figures 3F, 3G, and Table 1**). These results show that in both cases, the residence time is brief, and VASH interacts only transiently with microtubules. Based on the previously reported *K*_cat_ of VASH1 [33], which is 44.5/min (or 0.74/sec), the dwell time of both motion types only allows 1-2 catalytic cycles per interaction with the microtubule, which is consistent with our estimation of the number of detyrosinated sites in nascent detyrosinated puncta in super-resolution images of Nocodazole washout cells (**Figure 2F**). In addition, in our single molecule tracking videos, especially in cells with relatively high expression of VASH, the VASH trajectories appeared to overlap with most if not all microtubules (**Figure S4G**). We have also observed that the number of detyrosinated microtubules was much higher in VASH-EGFP overexpressing cells compared to untransfected cells (**Figure S5**). Additionally, Taxol treatment drastically increased both the kinetics of detyrosination and the percentage of detyrosinated microtubules (82 ± 25 % of total microtubule after 3 hr Taxol treatment, **Figure S6**), which is consistent with previous literature [3, 5, 51]. Overall, these results support that VASH can bind and interact with all microtubules. Further, we did not observe multiple VASH molecules binding to the same region consecutively, which is also consistent with the super-resolution data and suggests that microtubule binding cooperativity is likely not the dominant mechanism for detyrosination to build up continuously on a microtubule.

It is possible that VASH molecules may distinguish the detyrosinated versus tyrosinated α-tubulin tails on the microtubules, which may promote a preference for VASH to bind and dwell longer on microtubules with more tyrosinated tubulins to allow more catalytic cycles. To test this possibility, we carried out single molecule tracking experiments of VASH1 and VASH2 at 30 min post Nocodazole washout, when microtubules are largely reassembled but detyrosination is still mostly absent (**Figure 1**). We found that the percentage of Brownian and confined motions of either VASH1 or VASH2 were similar at 30-minute Nocodazole washout to the conditions where detyrosination has been established (**Figure S7, Table 1**). The residence time of Brownian (1.34 and 1.70 sec for VASH1 and VASH2) and confined (1.38 and 2.23 sec for VASH1 and VASH2) motions were also comparable and if anything, slightly shorter than those from cells that have more established detyrosination (**Figure S7, Table 1**). This suggests that VASH does not dwell longer on more tyrosinated microtubules in cells.

### A microtubule binding mutant of VASH2 exhibits reduced dwell-time on microtubules

Numerous correlative structural and functional experiments demonstrated specific mutations that separate VASH’s microtubule-binding from its free tubulin tail binding or its catalytic core. [3, 4, 32, 33, 44, 51–53]. To probe what factors can affect VASH’s residence time, we tested a previously reported VASH2 mutant (VASH2 4A) that is defective in microtubule-binding and has reduced detyrosination efficiency (**Figures 3A and 3F-G**) [51]. Single molecule tracking of this mutant showed that the defect in microtubule binding resulted in a significant decrease in the residence time of both confined and Brownian trajectories (**Figures 3F and 3G**) without changing the proportion of each motion type (**Figure 3E**). These results suggest that having a long enough dwell time on the microtubule is important for VASH’s ability to efficiently detyrosinate α-tubulin tails. Given that the decrease of bulk detyrosination level in V2 4A is much more pronounced than the decrease we observed in residence time (**Figures 3A and 3G**), it is possible that the on rate of this mutant is also lower compared to the WT. However, we cannot determine the association rate constant (k_on_) of VASH as this rate constant is concentration dependent and the concentrations of VASH in individual cells are unknown. Interestingly, we have also found that the VASH1 residence time is modestly lower compared to VASH2 in both confined and Brownian motions (**Figures 3F, 3G and S7, Table 1**), and that the relative detyrosination efficiency of VASH1 is also modestly lower than VASH2 (∼0.7 to 1, **Figure 3G**).

We further tested whether catalytic activity is crucial for VASH’s dwell time and interaction with microtubules. We tracked the single molecules of the catalytically inactive mutant VASH2 C158A (V2 C158A), which is virtually incapable of detyrosinating microtubules (**Figures 3A and 3G**) [51, 52]. The percentage of Brownian and confined motions of the V2 C158A was similar to WT VASH2 as were the residence time of Brownian and confined trajectories (**Figures 3E and 3F-G, Table 1**). These results suggest that VASH molecules do not change their mode of interaction with microtubules regardless of whether they catalyze detyrosination events.

### Simulations support that a detyrosination-stabilization feedback mechanism can generate a subpopulation of microtubules enriched with detyrosination

Collectively, both super-resolution microscopy and single molecule tracking experiments suggest that detyrosination is largely limited to only a few subunits at a time and occurs stochastically on microtubules. If detyrosination is established through stochastic and short-lived interactions between VASH enzymes and the microtubule network, it remains unclear how only a small subset of microtubules become enriched in detyrosination along their length. It has been shown that, through interaction with MAPs, detyrosinated microtubules become more stable. A previous in vitro reconstitution study showed that the level of microtubule dynamicity scales with the level of tyrosination [8]. Given that globally stabilizing microtubules in cells using Taxol accelerated the kinetics of detyrosination and increased the overall detyrosination levels (∼ 3-fold increase, **Figure S6**) [4, 5, 32], we hypothesized that the enrichment of detyrosination on a subset of microtubules starts from a stochastic process, but as a result of stochastic variations in the levels of detyrosination among individual microtubules, a subset of microtubules may become more stabilized than others. Consequently, this more stable sub-population of microtubules may accumulate more detyrosination, while the remaining microtubules turnover more readily and lose detyrosination. This feedback between detyrosination and stabilization can then lead to the appearance of aged, highly detyrosinated microtubules among young, tyrosinated microtubules.

Because the stability and detyrosination level of the microtubules cannot be readily determined and tracked simultaneously in live cells, we turned to simulations to assess whether a stochastic model with detyrosination-dependent microtubule stability can recapitulate experimentally observed detyrosination properties. Instead of mean-field kinetic models, which report only bulk microtubule dynamics [54, 55], we developed a computational model which additionally allows us to visualize the 3D spatiotemporal features of microtubule detyrosination by explicitly simulating the dynamics of individual microtubules, enzymes and the detyrosination states of the α-tubulin subunits. In our system (**Figure 4**), the microtubule (green) minus ends were stably embedded in the inner sphere representing the centrosome (pink) and the plus ends experience dynamic instability through Monte Carlo simulations [56, 57] within the outer spherical boundary representing the cell plasma membrane (green). Simultaneously, VASH molecules (small diffusing spheres) simulated via reactive Brownian dynamics, are capable of binding, unbinding and detyrosinating individual microtubules with certain probabilities estimated from experiments (**Figure 4** **and Supplementary Movie S8**).

**Figure 4:**
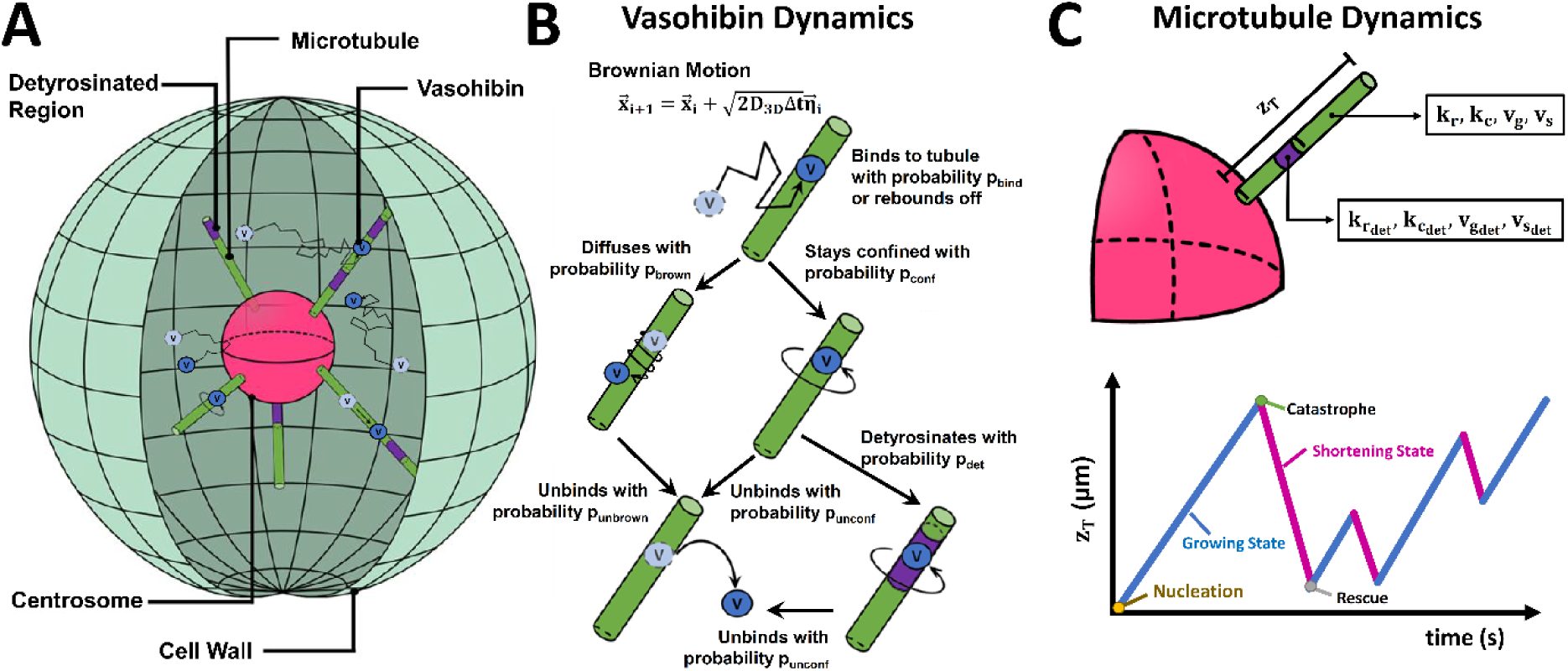
Schematics of our computational model and microtubule dynamic processes. **(A)** Simulated system. VASH enzymes (dark blue circles) diffusing inside the cell are capable to bind to existing microtubules (cylinders) and detyrosinate them (purple regions represent detyrosinated areas while green regions represent tyrosinated ones), locally increasing the stability of the microtubule. Light blue circles represent a previous position of a VASH enzyme in the trajectory (black arrow). **(B)** Enzyme dynamics. Upon binding to a microtubule (what happens with a probability p_bind_), a diffusing enzyme either becomes confined to an α-tubulin subunit with probability p_conf_, or it performs a random walk along the surface of the microtubule with a probability p_brown_. Confined enzymes and diffusing enzymes are capable of unbinding from a microtubule with probabilities p_unconf_ and p_unbrown_, respectively. Confined enzymes are also capable of detyrosinating the α-tubulin subunit they are bound to with probability p_det_. **(C)** Dynamic instability of a single microtubule. An α-tubulin subunit with growth and shortening rates v_g_ and v_s_, respectively, and rescue and catastrophe frequencies k_r_ and k_c_, respectively, changes its dynamic properties upon detyrosination (v_g_det__, v_s_det__, k_r_det__, and k_c_det__). A complete list of parameters are reported in **Table 2**.

**Table 2.**
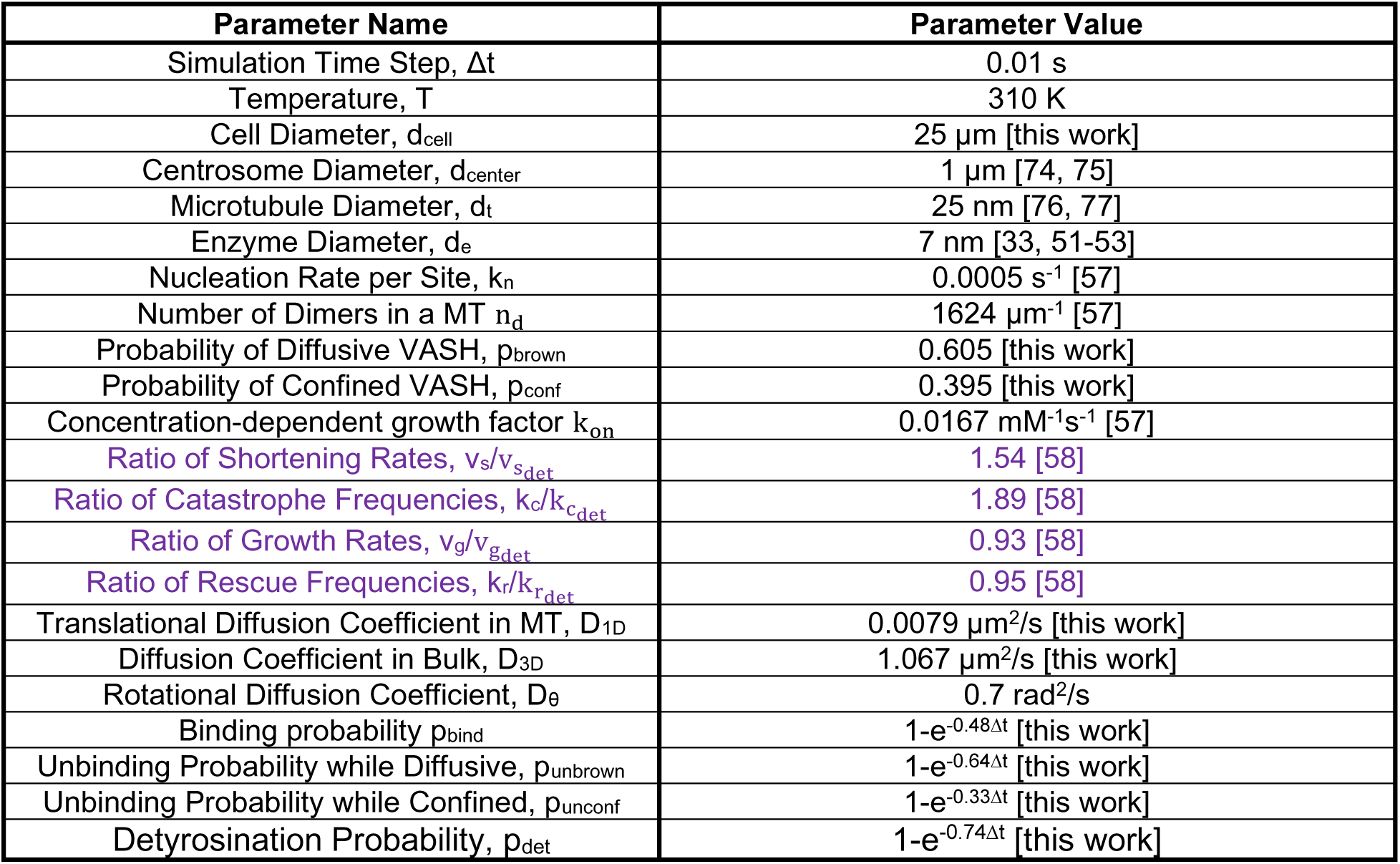
Common parameters used in simulations of the microtubule array. A timestep of A temporal resolution of 0.01 s was selected as an optimal value balancing total simulation time and accuracy. A rotational diffusion coefficient of 0.7 rad^2^/s was selected to randomize the point of escape of the bound enzyme on the microtubule. Binding, unbinding and detyrosination probabilities (p) are related to their associated times (T) by means of p = e^−Δt/T^.

### Parameters and assumptions of the simulations

Based on prior literature, our simulations assume the detyrosinated α-tubulins are stably present on the microtubule until it is released from the microtubule upon disassembly, and all nascently assembled microtubules, including the rescued segments are initially tyrosinated. Consequently, detyrosination level fluctuates as the microtubule plus ends experience dynamic instability.

The simulated probabilities of diffusive versus confined movement of VASH were based on the fractions of each motion class from our single molecule tracking data, and we assumed that VASH detyrosinated a subunit when confined on a microtubule, with a catalytic probability derived from the enzyme catalytic rate (0.74 s^-1^, [33]). The probability of VASH dissociation was based on the enzyme off-rate (1/residence time) and subsequently the binding probability was calculated based on the previously measured k_d_ (0.16 µM, [51]) between VASH and Taxol-stabilized microtubules. A complete list of parameters used in our simulations is shown in **Table 2.**

### Enrichment of detyrosination arises from stochasticity, high dynamicity of microtubules and detyrosination mediated microtubule stabilization

To the best of our knowledge, there is no established, quantitative model describing how microtubule stability evolves as detyrosination increases on a microtubule at the molecular level. We first assumed that stability is conferred at the individual subunit level. In this model (**Model 1**), the plus end dynamics transition to a more stable regime each time the plus end encounters detyrosinated subunits as it grows and shrinks. The more stable plus end has catastrophe and shortening rates reduced by a certain factor (∼65 % catastrophe frequency and ∼53 % shortening rate of the tyrosinated subunits), along with minor changes in rescue frequency and growth rate (see **Table 2**, purple letters). The percentage changes in the catastrophe and shortening were based on the ratios we obtained from a previous study that characterized these parameters for both dynamic and stable populations of microtubules in the same cell type [58].

To test how this model depends on the starting stability level of the microtubules, we simulated three different scenarios consisting of highly dynamic (set A), intermediately dynamic (set B) and fully static microtubules (set C, **Table 3, Figure S8**). To compare to our experimental data, we blurred our simulation resolution (0.6 nm) to the typical resolution of DNA-PAINT imaging (∼30 nm, about 50-fold lower resolution than the simulation) (**Figure S9, and Methods**). The cellular concentrations of VASH are unknown but are likely in the nanomolar range. We first studied the early stages of detyrosination establishment, similar to our nocodazole wash-out experiments. The buildup of detyrosination in the highly dynamic microtubules (set A) (**Figure S9**) most closely matched our experimental data, recapitulating the early kinetics of detyrosination. When the starting level of microtubule stability was higher (Set B and C), detyrosination accumulated faster and at higher levels, similar to our Taxol-stabilized microtubule experiments (**Figure S6**). We next evaluated whether detyrosination reaches a plateau once steady state is reached. We increased the enzyme concentration to reach steady state within a reasonable simulation time and found that detyrosination indeed plateaus at long times. Once again, as expected from our experiments using Taxol to stabilize microtubules (Figure S6), we found that higher initial microtubule stability led to faster detyrosination as well as higher plateau levels of detyrosination at steady state (Figure S9A).

**Table 3.** Parameter sets for microtubule plus-end dynamic instability.

| Parameter | Set A | Set B |
| --- | --- | --- |
| Initial Growth Rate $v_{g_0}$ (μm/s) | 0.142 | 0.192 |
| Shortening Rate $v_s$ (μm/s) | 0.188 | 0.218 |
| Catastrophe Frequency $k_c$ (s<sup>-1</sup>) | 0.053 | 0.026 |
| Rescue Frequency $k_r$ (s<sup>-1</sup>) | 0.086 | 0.175 |
| Tubulin Concentration (μM) | 25 | 35 |

Parameters were selected based on a previous study [57]. For **Set C**, microtubules were kept at their maximum lengths with their plus ends fixed at the cell boundary throughout the simulations.

We then sought to determine whether detyrosination-dependent stability affects detyrosination levels. Surprisingly, when we compared the buildup and steady state levels of detyrosination between **Model 1** and a control simulation (**control Model**) where microtubule stability does not change with the detyrosination state of the microtubule, we found little effect of the microtubule stabilization resulting from the detyrosination of individual α-subunits on the overall detyrosination levels, suggesting that the dynamicity of the microtubules overrules this stabilization (**Figure S10, set A**). We further took advantage of our spatiotemporal model to determine whether detyrosination becomes highly concentrated on a subpopulation of microtubules under **Model 1**. We used the distribution of the percentage of detyrosination per microtubule to assess the enrichment of detyrosination. If detyrosination becomes enriched on a subset of microtubules, we would expect a bimodal distribution with a population having low and a population having high detyrosination levels. In both the **control Model** and **Model 1**, the percent detyrosination histograms of highly dynamic microtubules showed a broad distribution, which did not fit to a double-Gaussian distribution (**Figure 5A**), suggesting little or no detyrosination enrichment on microtubule sub-populations. Hence, while **Model 1** captured the early kinetics of detyrosination buildup, it did not differ drastically from the **control Model** and did not recapitulate the enrichment of detyrosination on a subset of microtubules. We reasoned that, under **Model 1**, the stability conferred by detyrosination is limited by the location of the detyrosinated subunit on a microtubule, and only the few subunits clustered near the microtubule plus end will be able to provide the stability required to slow down depolymerization. Subunits placed far from the plus end will not confer any stability. Alternatively, it is possible that there is a threshold level of detyrosination that needs to accumulate for a microtubule to have increased stability, regardless of the spatial distribution of detyrosinated subunits along the microtubule lattice. To test this hypothesis, we established **Model 2**, in which a microtubule only becomes more stable (**Table 2**) when it reaches a certain threshold of overall detyrosination. In this model, the evolution and steady state levels of detyrosination did drastically differ from the **control Model** at a range of microtubule stability thresholds (**Figure S10B**).

**Figure 5:**
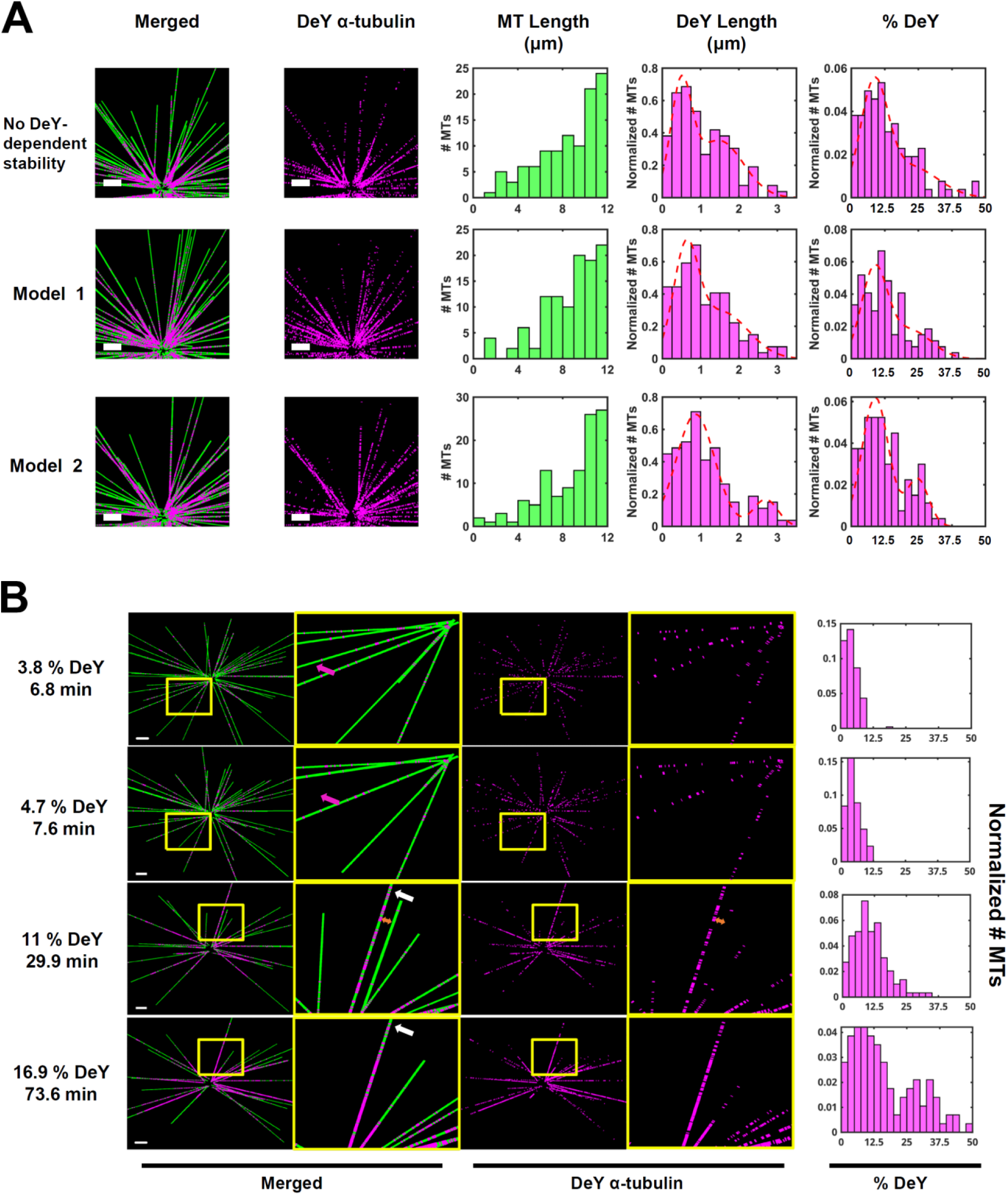
Due to their increased stability, highly detyrosinated microtubules co-exist with newly polymerized ones. **(A) (LEFT)** Representative images showing zoomed regions of highly dynamic (set A) microtubules at total detyrosination level of 12% with 500 nM VASH. A VASH concentration higher than the one considered in **Figure S9** was used to achieve detyrosination levels of 12% for our dynamic microtubules in manageable simulation times. Detyrosinated α-tubulin subunits are shown in magenta and β-tubulin in green. Scale bars, 2 µm. Images were blurred to experimental resolution (Methods). **(RIGHT)** Histograms of the total microtubule length, detyrosinated segment length, and percentage of detyrosination. Fully depolymerized microtubules were neglected in our histograms. Percentages of detyrosination were fitted by double-Gaussian distributions (red dashed lines), and the fit parameters are listed in **Supplementary Table 1**. **(B) (LEFT)** Images showing a region of randomly selected 50 out of 250 microtubules (simulated using Model 2) in the presence of 500 nM VASH at time points and total detyrosination levels indicated. Detyrosinated α-tubulin subunits are shown in magenta and β-tubulin in green. Scale bars: 2.5 μm. Yellow open boxes indicate regions selected for zoom. Images were blurred to experimental resolution. Note that a tyrosinated microtubule (magenta arrows) rapidly shortened (0.8-minute lifetime) while a highly detyrosinated one persisted for over 45 minutes (white arrows). The orange bi-direction arrow shows an example of a highly detyrosinated microtubule positioned right next to a tyrosinated one at 29.9 min. Note that they mostly tyrosinated microtubule had already disassembled by 73.6 min. **(RIGHT)** Histograms of the percentage of detyrosination of the total 250 microtubules at the indicated time points.

When the threshold in **Model 2** was set to 20%, meaning that when the detyrosination level of a microtubule reaches 20% of its maximum possible length at microscopy resolution, this individual microtubule will become more stable overall. Under this hypothesis, two distinct populations of microtubules emerged based on their detyrosination percentage and length. These two populations could be fit to two well-separated Gaussian peaks (red dashed lines, **Figure 5A**, 3^rd^ row, **Table S1**), suggesting that a small subpopulation of microtubules had substantially higher detyrosination levels than the rest, which was also evident visually (**Figure 5A-B**). Such well-defined bimodality was not observed when the threshold for microtubule stabilization was set to 5 or 10 % detyrosination levels (**Figure S11**), where microtubules gain stability more rapidly. Similarly, we did not observe this bimodality under **Model 1** with microtubules that are more inherently stable (set B and C, **Figure S11**). In fact, the detyrosination percentage distributions from more stable microtubules were much less heterogeneous than those from dynamic microtubules. This heterogeneity resulting from microtubule dynamicity likely promotes the emergence of detyrosination enrichment on microtubule subpopulations. Indeed, a close examination of the progression of the detyrosination on highly dynamic microtubules under **Model 2** (with 20 % threshold) showed that relatively high level of detyrosination along the entire length of some microtubules slowed down depolymerization allowing these microtubules to further accumulate detyrosination. In the region shown in **Figure 5B**, a sparsely detyrosinated microtubule lasted only 0.8 minute (**Figure 5B**, magenta arrows), whereas a highly detyrosinated microtubule persisted for over 45 minutes (**Figure 5B**, white arrows). Particularly, we observed a significant amount of detyrosination accumulation on this microtubule between 29.9 min and 73.6 min (**Figure 5B**, white arrows), giving rise to the phenomenon of aged, highly detyrosinated microtubules co-existing with young tyrosinated ones (**Figure 5B**, bi-directional arrow). **Model 2** also provided a better match to the experimental results for the early kinetics of detyrosination buildup than **Model 1** (compare **Figure S9** and **Figure S12**).

Altogether, our simulations suggest that the enrichment of detyrosination on a sub-population of microtubules requires two components (**Figure 6**): (1) highly dynamic microtubules that lead to the emergence of detyrosination heterogeneity via a stochastic detyrosination process, and (2) a threshold-based microtubule stability mechanism that leads to the enrichment of detyrosination on a subset of microtubules that accumulate a certain level of detyrosination. When microtubules are more stable (e.g. through Taxol stabilization), the initial heterogeneity in microtubule detyrosination as well as the feedback from detyrosination induced stabilization are lost such that detyrosination is no longer enriched on a subpopulation of microtubules

**Figure 6:**
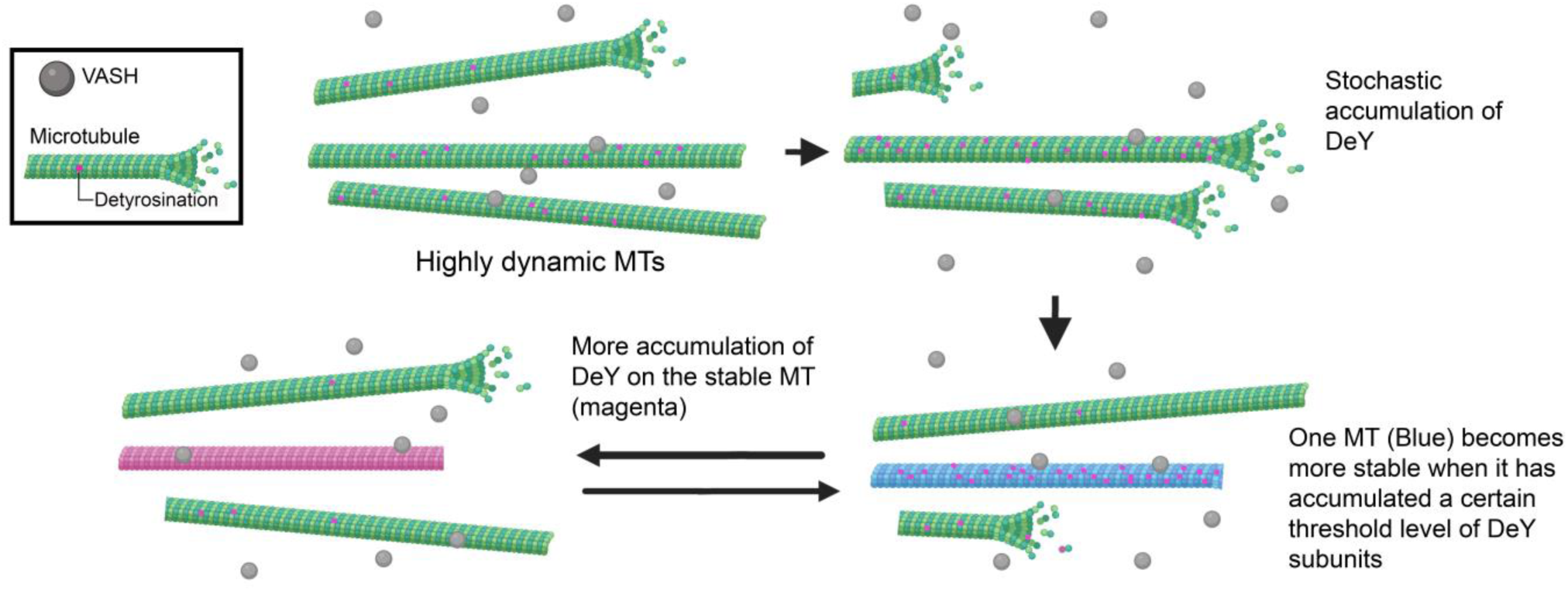
The feedback between microtubule stability and detyrosination. In the cells where MAPs are present, VASH stochastically acts on microtubules and by chance detyrosination level varies among different microtubules. Over time, some microtubules accumulate more detyrosination above a certain threshold and become stable (shown in light blue), while the rest turnover readily and lose detyrosination. The stability of the microtubule allows more accumulation of detyrosination events (exemplified by a purple microtubule here) and eventually lead to much higher stability when detyrosination reaches an even higher threshold. The stability and detyrosination form a feedback loop, indicated by the anti-parallel arrows. It is possible that additional factors can stabilize microtubules regardless of microtubule detyrosination level, which will accelerate the evolution of stability, also leading to accumulation of more detyrosination on stable microtubules.

## Discussion

Numerous PTMs are found in selected subsets of microtubules in cells, and how microtubules are segregated into these functional subpopulations has been a long-standing question [2]. Detyrosination is among the first microtubule PTMs [59, 60] that were identified and, unlike other PTMs such as acetylation, detyrosination can be highly compartmentalized to bias organelle transport mediated by motors [16, 31]. Using a combination of quantitative super-resolution, live cell single molecule tracking and spatiotemporal stochastic simulations, we propose that detyrosination is concentrated on a small subset of microtubules through a stabilization-detyrosination feedback mechanism starting from a stochastic process. Early after nocodazole washout, the quantitative DNA-PAINT super-resolution images provided us the insight that detyrosination occurs primarily on one α-tubulin tail at a time (∼60% of all initial sites contain a single detyrosinated subunit) and these initial detyrosinated sites are many tubulin subunits away from each other (median distance, 179 nm) (**Figure 2**). Regardless of the type of enzyme that is at play, these results suggest that consecutive or cooperative detyrosination on the microtubules is not prevalent, instead, detyrosination is sporadic. Furthermore, throughout the time course post washout, the proportion of single detyrosinated sites decreased but persisted and these sites appeared closer to each other, suggesting that continuous detyrosination in untreated cells results from the merging of separate, single detyrosination events as they appear more frequently with time.

We tracked VASH as the primary detyrosinase, which contributes to the majority of total detyrosination in many cell lines and in the mouse brain [3–5]. These data showed that the brief interaction of VASH with the microtubule only allows 1-2 catalytical cycles for VASH to detyrosinate, which is consistent with the results from our super-resolution quantification. In all our experiments, VASH1 and VASH2 do not show major differences except that VASH2 expression results in modestly higher detyrosination than VASH1 (**Figure 3A**), and accordingly, VASH2 consistently has a slightly longer dwell time in all motion categories (**Figures 3F and 3G**, Table1). These results are consistent with previous literature showing conserved structures between these two isoforms. These results are also consistent with a recent in vitro study, which used total internal reflection fluorescence microscopy to track the binding of VASH1 and VASH2 to microtubules [45]. We note that in the in vitro work VASH2 residence time on the microtubule was dramatically longer than VASH1 residence time [45]. It is possible that in the cell context, presence of other microtubule associated proteins (MAPs) and PTMs dampens the dramatic differences in the binding mode of these two enzymes, as we found that the residence time in cells differ only slightly. Interestingly, about half of the total trajectories are diffusive on microtubules, and given the slow catalytic rate of VASH (0.74 s^-1^ for VASH1 [33]) and the limited number of detyrosination events found in detyrosination puncta in our super-resolution images, these diffusive events that span many tubulin subunits within 1-2 seconds are unlikely the detyrosinating events. We interpret that VASH, through the diffusion, surveys the microtubule surface to locate a substrate, as we have observed a small fraction of trajectories that transition from diffusive behavior to confined behavior (**Figure S4E** and **Supplementary Movie S7**). Such “facilitated 1D diffusion” mechanism has previously been suggested to help transcription factors find their binding sites within chromatin more rapidly compared to 3D diffusion [61, 62]. Similarly, diffusion along the microtubule can potentially help reduce VASH’s search time for a target site. So far, it is unclear what governs VASH to be confined as opposed to diffusing along the microtubule lattice. We have found no evidence suggesting that VASH changes its behavior if the α-tubulin tail is already detyrosinated, because we observed similar proportion of diffusive and confined motions when we tracked VASH early after nocodazole washout, when microtubules are mostly tyrosinated. These results are consistent with in vitro work showing very little differences in the interaction of VASH with tyrosinated and detyrosinated microtubules [45].

It is possible that MAPs present on microtubules may influence whether VASH can be stably bound to one α-tubulin tail for the amount of time the enzyme requires for detyrosination. For example, a recent study suggested that presence of MAP4, but not phosphorylated MAP4, on microtubules may block the local access of VASH2 in cardiomyocytes [63]. Previous cryo-EM studies showed that VASH, presumably stably bound to microtubules, engages with two α-tubulins from two adjacent protofilaments [32, 45], and structural studies also suggested that the C-terminal tail of α-tubulin is supposedly occupied by the catalytic pocket of VASH for detyrosination to occur [32, 33, 51, 52].

In our simulations, we constructed an idealized system incorporating the two main mechanisms, the stochastic detyrosination and the stabilization of microtubules by detyrosination. While similar approaches have been employed to characterize diverse biological processes involving molecular recognition, such as the search of regulatory proteins for specific DNA-binding targets [64, 65] and the clustering of chemotaxis sensing complexes [66, 67], to the best of our knowledge, this is the first implementation of reaction-diffusion algorithms to characterize how dynamic microtubules are modified by an enzyme over time with 3D spatial resolution. Our simulations captured the nascent detyrosination kinetics shown by our experiments and showed that the levels and kinetics of detyrosination are dependent on microtubule stability and enzyme concentration. We also observed the appearance of “old” microtubules highly enriched in detyrosination next to “young” tyrosinated microtubules. These results support that in the stochastic system, the dynamicity of the tyrosinated microtubules and the stability feedback mechanism together promote the enriched detyrosination (**Figures 5 and 6**).

It is unclear how microtubule stability precisely evolves as detyrosination progresses on microtubules in the cells. In our simulations, we have found that a threshold-based model (**Model 2**) led to a much more pronounced detyrosination enrichment on a subset of microtubules. This is possibly because this model is independent of spatial distribution of detyrosination on a microtubule whereas in **Model 1**, microtubule stability changes most significantly when detyrosinated tubulin are clustered near the plus end. We also observed partially detyrosinated microtubules in all simulations, which were not observed experimentally in epithelial cells. This discrepancy could be due to a couple of reasons. First, it is possible that we underestimated the stability of detyrosinated microtubules. Indeed, in either **Model 1** or **Model 2**, the change in stability upon detyrosination is only a factor of ∼2 in reduction of catastrophe frequency and shortening rate, which produced limited effect on overall detyrosination level. Second, the microtubule stability increase is likely more complex in cells than what our models assume. In **Model 1**, when stability change is conferred at single tubulin level, the overall stability will continue to rise with detyrosination increase, but a microtubule with a detyrosinated subunit close to the plus end will not depolymerize as readily as a microtubule with a detyrosinated subunit close to the minus end, despite having similar levels of detyrosination. In **Model 2**, the microtubule stability goes through a binary switch rather than a continuous or step-wise increase as the detyrosination builds up, which limits the lifespan of the highly detyrosinated microtubules that may otherwise persist even longer [8, 25, 26, 68]. Additionally, in our simulations the detyrosination levels on the enriched microtubule sub-population did not reach 100% even after resolution blurring, regardless of the simulation model. This discrepancy between the simulations and experimental results is likely due to the fact that our segmentation algorithms may miss small gaps in detyrosination that exist along the microtubule lattice, leading to an artificially higher level of detyrosination. It is possible and likely that the entire lattice is not fully saturated with detyrosinated subunits.

It has been observed that unlike in epithelial cells or neurons, some other cell types do not have distinct detyrosinated versus tyrosinated microtubule populations, but rather partial detyrosination on many microtubules like in our simulations [19]. Decades of literature have provided highly variable, cell type-dependent measurements for the catastrophe and shortening rates of microtubules. This is likely due to various factors, including concentration of VASH and MAPs, and the composition of MAPs, which likely account for the cell type-dependent degree of stability. These variables may impact how restricted or widespread detyrosination is depending on the cell type. Therefore, simulating detyrosination process with additional models for microtubule stabilization may generate testable predictions for future experiments.

It is interesting to postulate whether this feedback mechanism can explain enrichment of other PTMs on subsets of microtubules, for example, stable microtubules are also enriched in acetylation and glutamylation [1, 2]. Incorporating the results and simulations from our study, we reason that the microtubules which persist longer will allow the time-dependent accumulation of PTMs if polymers are the preferred substrate and the enzyme acts on a time scale that is much slower than the inherent stable periods of the microtubules. Additionally, we found that the enzyme concentration can strongly influence the outcome when the enzyme is diffusion limited (**Figure S10A**). In our study and a previous report [43], L40 acetylation is re-established in the cells much faster than detyrosination after nocodazole washout and acetylation is also much more prominent than detyrosination, occurring on 70% of all microtubules. Previous in vitro work indeed showed that acetylation, similar to what we find for detyrosination in cells, is stochastic and that TAT can access and enter the microtubule lumen not only through microtubule ends but also within the microtubule lattice [34]. These results are consistent with the cellular patterns of widespread acetylation and suggest that two different stochastically acting enzymes can give rise to differential modification kinetics and different levels of modified microtubules, likely by tuning the enzyme concentration, the enzyme catalytic rate and the enzyme binding time to the microtubule with respect to the timescale of microtubule stability. It is also possible that since acetylation happens in the microtubule lumen, in cells, TAT becomes trapped inside the microtubule and that this confinement helps build continuous stretches of acetylation [43]. It has been shown that glutamylation is more efficient on the microtubule polymer as well (Audebert et al., 1993), however, the relationship between glutamylation and stability is complex, for example, polyglutamylation regulates microtubule severing mediated by spastin and katanin in a biphasic manner, therefore at an intermediate level, polyglutamylation results in less stable microtubules whereas high level polyglutamylation stabilizes microtubules [69–73]. Since tubulin PTMs play diverse roles and that the modifying enzymes have different catalytic rates (reviewed in [2]), the proposed feedback mechanism between PTMs and microtubule stability in our study provides a general framework for determining the kinetics, steady-state levels and patterns of modified microtubules in different cell types. Our experimental and computational approach should provide useful tools in studying how various PTMs are established in the cellular context.

### Captions for videos

**Movie S1** A movie showing Brownian events (a and b in Fig 3C) from two VASH2-Halo molecules, pseudo-colored in yellow on microtubules (magenta) labeled by Tubulin Tracker. Scale bar, 1 µm. The movie is playing at 1× speed or 20 frames per second (fps).

**Movie S2** A movie showing a VASH1-Halo molecule (Fig S4C) diffusing back and forth on the microtubules. Scale bar, 1 µm. The movie is playing at 1× speed or 20 fps.

**Movie S3** A movie showing a prolonged diffusion from a VASH1-Halo molecule (Fig S4D) on the microtubule dense area. Scale bar, 1 µm. The movie is playing at 1× speed or 20 fps.

**Movie S4** A movie showing a prolonged diffusion from a VASH2-Halo molecule (Fig S4D) on the microtubule-dense area where microtubules possibly bundle or criss-cross. Scale bar, 1 µm. The movie is playing at 1× speed or 20 fps.

**Movie S5** A movie corresponding to Fig 3C, showing a VASH2-Halo (JF549) molecule is confined on Tubulin Tracker labeled microtubules (magenta). Scale bar, 1 µm. The movie is playing at 1× speed or 20 fps.

**Movie S6** A movie showing a VASH1-Halo molecule is confined on a microtubule (Fig S4C). Scale bar, 1 µm. The movie is playing at 1× speed or 20 fps.

**Movie S7** An example movie showing a VASH2-Halo molecule (Fig S4E) transition between Brownian and confined motions while staying on microtubules. Scale bar, 1 µm. The movie is playing at 1× speed or 20 fps.

**Movie S8.** An example movie showing a system of 10 microtubules (cylinders) and 10 VASH molecules (spheres) in a cell of radius 5 μm. Microtubule and enzyme diameters were increased for visualization purposes as well as the size of an α-subunit. Tyrosinated regions of the microtubules appear in green while detyrosinated regions appear in purple. Multiple parameters were modified to show faster detyrosination. As an example, one VASH molecule (black) was tracked during the simulation time and its cumulative trajectory is shown in the lower left panel.

## Methods

### Cell culture expression constructs

BSC-1 or U2OS cells (BS-C-1; CCL-26; American Type Culture Collection) were routinely maintained in MEM or DMEM media (ThermoFisher) supplemented with 10% (vol/vol) FBS, 1x Glutamax, 1x Anti-Anti, 1 mM sodium pyruvate at 37 °C with 5% CO_2_. U2OS and 293T cells were cultured in DMEM media supplemented with 10% (vol/vol) FBS, 1x Glutamax, 1x Anti-Anti, 1 mM sodium pyruvate at 37 °C with 5% CO_2_. All culture media and supplements were purchased from ThermoFisher.

### DNA constructs and lentivirus production of VASH and SVBP coexpression

For transient expression from plasmids, VASH1-eGFP or VASH2-eGFP [51] (kind gifts from Dr. Marie-Jo Moutin) were each co-expressed with SVBP-Myc-FLAG. Briefly, BSC-1 cells were grown to 60% confluency in 6-well plate and were co-transfected with 500 ng each of VASH-eGFP and SVBP-Myc-FLAG using Lipofectamine 2000 (ThermoFisher) following manufacturer’s instructions. The media was replaced the next day and the cells were trypsinized and re-seeded onto Lab-TekII imaging chamber (Nalge Nunc, ThermoFisher). The transfected cells were imaged ∼40 h post transfection.

Lentiviral expression constructs pLV-CBh-SVBP-Myc-FLAG:IRES:VASH-HALO, co-expressing SVBP and VASH WT or mutants, were commercially made (VectorBuilder, Chicago, IL and Genscript). Briefly, human SVBP-Myc-FLAG sequence were synthesized and inserted downstream of a CBh promoter and VASH coding sequences were placed downstream of an IRES element. Human VASH1 WT, VASH2 WT, VASH2 C158A, VASH2 4A coding sequences were amplified from previously published mammalian expression constructs VASH1-eGFP, VASH2-WT-eGFP, VASH2 C158A-eGFP, VASH2-4A-eGFP [51] (kind gifts from Dr. Marie-Jo Moutin).

Lentiviral particles were packaged produced by co-transfecting the expression construct pLV-CBh-SVBP-Myc-FLAG:IRES:VASH-HALO and packaging constructs (kind gifts from Dr. Ben Prosser) pCMV-VSVG, pRSV-REV, pCgpV at 3:1:1:1 ratio into 293T cells. Briefly, 45 µg of expression construct and 15 µg of each of the packaging constructs were mixed in 900 µl Opti-Mem media (ThermoFisher) and subsequently mixed with 900 µl Opti-Mem media containing 90 µl Lipofectamin2000 for 5 min. The mixture was then added to a 10 cm-plate of T293 cells at 50-60% confluency for overnight incubation at 37 °C. The media was replaced with fresh growth media the next day and the cells were allowed to produce virus for the next 48 hr. The media containing the packaged viral particles were collected and spun at 100 ×g for 5 min and the supernatant was passed through a 0.45 µm filter, aliquoted and stored at -80 °C.

### Nocodazole washout and immunostaining for DNA-PAINT

BSC-1 cells were grown in Lab-TekII imaging chamber. For cells treated with Nocodazole, the BSC-1 growth media containing 33 µM Nocodazole was filtered through a 0.45 µm filter and added to the cells for three hours at 37°C. For cells that are allowed to reassemble microtubule after Nocodazole treatment (washout conditions), the cells were then washed three times with warm complete growth media and then incubated in growth media for different amount of time as indicated in the results section. For washout experiments followed by Taxol treatment, cells were quickly washed three times after Nocodazole treatment and was incubated with 2 µM Taxol or the equivalent amount of DMSO (0.04%, vol/vol, labeled as untreated) for 30 min at 37°C. To fix the cells, they were briefly (10-30 s) permeabilized by incubating with extraction buffer (80 mM PIPES-KOH pH 7.1, 1 mM EGTA, 1 mM MgCl_2,_ 0.5% Triton X-100 (vol/vol), 10% glycerol (vol/vol)) warmed to 37°C and immediately replaced by the fixation buffer (3% paraformaldehyde (PFA) and 0.1% glutaraldehyde in PBS) at 37°C for 10 min. Cells were then washed twice with PBS, incubated with fresh 0.1% NaBH4 (wt/vol) solution for 7 min and washed again three times with PBS. The fixed cells were then permeabilized for 10 minutes with 0.2% Triton X-100 (vol/vol) in PBS and blocked for one hour at 25°C using Primary Blocking Buffer (10% donkey serum (vol/vol), 0.2% Triton X-100 (vol/vol), and 0.05 mg/mL sonicated salmon sperm single stranded DNA (Stratagene, La Jolla, California) in PBS). Salmon sperm DNA was denatured at 95 °C for 3 minutes before addition to the buffer and was included in the blocking buffer to reduce off-target binding of DNA-PAINT secondary antibodies and imagers to nuclear DNA. Cells were then incubated with primary antibodies at 1:100 unless otherwise indicated in the figure legends in Primary Blocking Buffer, for one hour at 25°C or at 4°C overnight. Following primary antibody incubation, cells were washed once with PBS, followed by three five-minute washes with Wash Buffer (Massive Photonics, Gräfelfing Germany). Cells were then incubated with secondary antibodies (anti-mouse and anti-rabbit DNA-PAINT secondary antibodies, Massive Photonics) at 1:100 dilution in Antibody Incubation Buffer (Massive Photonics). Cells were then washed once with PBS briefly, followed by three ten-minute washes with 1X Wash buffer (Massive Photonics). Finally, cells were washed once for five minutes with PBS, and stored in fresh PBS at 4°C until imaging.

### DNA-PAINT Imaging

DNA-PAINT images were acquired using a Nanoimager (ONI, Oxford, UK) equipped with 405-, 488-, 561- and 640-nm lasers, 498-551- and 576-620-nm band-pass filters in channel 1, 666-705-nm band-pass filters in channel 2, a 100 x 1.45 NA oil immersion objective (Olympus), and a Hamamatsu Flash 4 V3 sCMOS camera. All images were acquired with 100 ms exposure at 30°C using HILO illumination [78]. Before imaging, imager DNA oligos (Massive Photonics) complementary to the conjugated secondary antibody docking oligos were diluted in Imaging Buffer (Massive Photonics) and added to imaging wells. To perform two-color imaging, for beta tubulin and acetylated tubulin, 0.25 nM of Imager-1 Cy3B or Imager-1 ATTO655 was used, and 5,000 total frames were acquired. This was sufficient to capture a continuous image of microtubules, allowing for their use as a location reference for detyrosination signal in subsequent analysis. For detyrosinated tubulin, 0.5 nM of Imager-2 Cy3B or Imager-2 ATTO655 was used, and 10,000 total frames were acquired. The laser program (Oxford Nanoimaging) alternates acquisitions between the two channels (100 frames of beta or acetylated tubulin followed by 200 frames of the detyrosinated tubulin) until the total 15,000 frames were recorded. For single-color imaging of acetylated microtubules for Nocodazole washout experiments, 0.5 nM of Imager-1 Cy3B or Imager-1 ATTO655 was used were used and 10,000 total frames were acquired. Images were then drift-corrected and filtered using the NimOS localization software (ONI) with filtering parameters as follows: X/Y localization precision (0-30 nm), Sigma X/Sigma Y (10-150 nm). All images regardless of experimental condition were filtered using these same parameters.

### Western blot

U2OS cells were grown to 50-60% confluency in 6-well plate and incubated with 400 µl lentivirus carrying the appropriate expression cassettes in a total volume of 1 mL culture media containing final concentration of Polybrene at 10 µg/mL per well overnight. The lentivirus containing media was removed the next day and the cells were trypsinized and seeded onto new wells to continue growth for another day. The cells were trypsinized and pelleted 40 hr post viral transduction and the pellets were solubilized with 1x LDS Sample buffer (ThermoFisher) supplemented with 1x Sample Reducing agent (ThermoFisher) and boiled for 7 min at 98 °C. The samples were resolved on a 10% Bis-Tris gel (Novex) and transferred to nitrocellulose blot. The blot was incubated with Odysee TBS blocking buffer (Licor) for 1 hr at 25 °C followed by incubation with primary antibodies, anti-Halo (1:1000, mouse, Promega), anti-Myc (1:100, Rabbit, Proteintech), anti-detyrosinated α-tubulin (1:1000, Rabbit, Abcam) and anti-GAPDH (1:2000, mouse, Proteintech) overnight, and the blot washed three times with 1xTBST (ThermoFisher) and secondary antibodies (680-anti mouse, 800-anti rabbit) at 1:10000 for 1 hr at 25 °C followed by three washes of TBST. The Western blot was imaged on Licor imaging system and the signal was quantified using the Licor imaging studio software. Detyrosination level was normalized to GAPDH level. To compare the exogenous level of detyrosination due to the ectopic expression of VASH-Halo, detyrosination was first quantified and normalized to GAPDH, and the basal level of detyrosination in un-transduced U2OS was then subtracted from all conditions that express VASH-Halo. The subtracted detyrosination level was then normalized to VASH2-Halo level. Detyrosination level due to VASH2 WT expression was arbitrarily set to one and all other conditions were described as a relative detyrosination level normalized to VASH2 WT (Figure 3A and 3G).

### Single molecule imaging of VASH-Halo

U2OS cells were grown to 50-60% confluency in a 12-well plate and transduced with 100 µl of lentivirus carrying the appropriate expression cassettes in a total volume of 400 µl culture media containing final concentration of Polybrene at 10 µg/mL per well overnight. The lentivirus containing media was removed the next day and the cells were trypsinized and seeded onto Lab-TekII imaging chamber and allowed to grow overnight. On the day of imaging (∼40 hr post viral transduction), cells were incubated with 20 nM of Halo ligand JF549 (Promega) in growth media for 15 min at 37 °C followed by three washes of warm growth media. Then the cells were labeled with Tubulin Tracker Deep red (ThermoFisher) following the manufacturer’s instruction. Briefly, the cells were incubated with 1× Tubulin tracker (1 µM) in growth media for 30 min at 37 °C. The cells were then washed three times with imaging media (L-15 lebovitz media (ThermoFisher) supplemented with 10% (vol/vol) FBS, 1× Glutamax, 1× Anti-Anti, 10 mM HEPES, 770 µg/mL Probenecid) and left in imaging media for imaging for no more than two hours. For single molecule imaging post Nocodazole washout, the cells were grown, infected with lentivirus and seed similarly as aforementioned. Prior to imaging, the cells were treated with 33 µM Nocodazole (0.45 µm membrane filtered) for 3 h at 37 °C. The cells were then quickly washed three times with complete DMEM culture media and incubated with Tubulin Tracker Deep red for 30 min at 37 °C, followed by similar wash and imaging steps with imaging media.

Single molecules and microtubules are visualized and recorded using an Nanoimager (ONI) warmed to 30 °C. For each recording, the microtubule confocal image was first captured in the confocal mode on Nanoimager with 500 ms exposure per slice using 640 nm laser at 65%-75% power for 10 consecutive slices. Immediately after, VASH-Halo was imaged in the widefield mode with HILO illumination using 560 nm laser at 15% laser power and the images were acquired at 50 ms exposure for 2500 frames, or a total of 125 s.

### Data Analysis

#### DNA PAINT

DNA PAINT images were analyzed using a custom-written Matlab code modified from a previous version [16]. Drift-corrected and sigma filtered (Nanoimager software, Oxford Nanoimaging) localizations were Voronoi segmented and clustered based on consistent Voronoi area threshold across all conditions (616.005 nm^2^ for detyrosinated tubulin and 5475.6 nm^2^ for beta/acetylated tubulin, each with minimum of 5 localizations per cluster/puncta). The large Voronoi area threshold for beta/acetylated tubulin is to ensure microtubules were clustered as a continuous network for localization references. Detyrosinated puncta that are denser (in localization per Voronoi area) than two standard-deviation from the mean were excluded from the detyrosinated images as non-specific background noise. Microtubule network clusters with Voronoi area less than 41067 nm^2^ were discarded as background noise. Detyrosinated microtubules were then overlaid with the reference microtubules and the detyrosinated puncta with ≥40% of their area overlapped with microtubules were considered colocalized with microtubules. The microtubule-colocalized detyrosinated puncta areas were calculated from the bounding puncta areas contoured by a Matlab built-in alphaShape function. Colocalized detyrosinated puncta from all experiments were filtered against a minimum area of 400 nm^2^ (area calculated from alphaShape objects), which is considered the resolution limit for DNA-PAINT. The lengths of the detyrosinated puncta were defined as the longest axes from principle component analysis (PCA) rotation of the puncta. The distance to the nearest punctum was the smallest Euclidean distance between borders of puncta.

To calculate the contribution of background noise to all detyrosinated signal, the puncta that have density (number of localizations per Voronoi area) larger than two standard-deviation away from the mean were first excluded. Then detyrosinated image was flipped horizontally and then vertically with respect to the microtubule reference image. The fraction of false positive co-localization was determined as the sum of the area of the flipped detyrosinated α-tubulin image co-localized with the β-tubulin image divided by the sum of the area of the original detyrosinated α-tubulin image co-localized with the β-tubulin image.

To calculate the number of detyrosination site per puncta, we used the detyrosination image obtained by staining with anti detyrosinated α-tubulin antibody at 1:2500 dilution as the single detyrosination calibration. The detyrosinated images (Figure 2D, 2E and Figure S2) were post-processed and colocalized with acetylated microtubules same as described above. The number of localizations per colocalized detyrosinated punctum was plotted as a histogram distribution and fitted to a lognormal function:

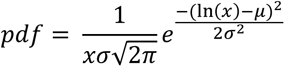

The x is the number of localizations. The mean (*µ*) and variance (*σ*) were obtained from the fit. The 2-, 3-, 4-, 5-detyrosination site calibration functions were obtained by linear convolution of the single detyrosination site calibration function from the lognormal fit using a custom-written Matlab code. The number of localizations per colocalized detyrosinated puncta from images at 30 min, 60 min and 90 min post Nocodazole washout were plotted as histograms and were fitted to the combination of calibration functions with weights w_1_, w_2_, w_3_, w_4_, w_5_ corresponding to the proportions of 1, 2, 3, 4, 5 detyrosination sites. The 3+ sites in the graph of 90 min post Nocodazole washout (Figure 2F) corresponds to the combined weight of w_3_, w_4_ and w_5_.

The total experimental detyrosination level (Figure 5B) is the calculated as the area sum of the detyrosinated clusters were divided by the area sum of the total microtubules for each cell.

### Single molecule tracking and analysis

Single molecules of VASH-Halo were tracked with ImageJ (NIH) plugin Trackmate [79]. Spots were detected using the LoG detector with 0.8 µm diameter threshold and subpixel localization and were subsequently linked as trajectories using Simple Lap Tracker with 0.6 µm maximum linking distance and maximum 1 frame gap. The linked trajectories were exported as XML files and analyzed using a custom-written MATLAB code. Briefly, Mean square displacement (MSD) of each trajectory is fitted to the power law function as below [48]:

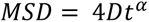

The D is the diffusion coefficient, t is the time lag between time points of the trajectories, and the first 25% of time lag of the MSD was fitted to extract the α exponent. Only trajectories ≥ 10 frames (0.5 s) were analyzed. Trajectories were then overlaid with the corresponding microtubule confocal image which were binarized using the local thresholding function in ImageJ. Trajectories with less than 20% of their segments overlapped with microtubules were considered not on the microtubule network. The 20% overlap threshold accounts for certain degree of microtubule fluctuations during the 125 s imaging time at 30 °C (Fig S4A). All trajectories were then manually inspected against the original confocal images to correct any mis-categorization. The motions were then classified based on the value of the α exponent as follows, α ≤ 0.8, confined; 0.8 < α < 1.5, Brownian; α ≥ 1.5, directed, and trajectories in all categories are filtered with coefficient of determination R^2^ ≥ 0.8.

### Residence time calculation

The residence time was calculated for Brownian and confined categories separately. In each category, all trajectories were pooled and plotted as survival fraction of molecules using 1-Cumulative Distribution Function of the trajectory duration. We then fit the survival curve with a two exponential decay function as follows ([49, 50, 80].

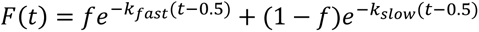

The *f* is the fraction of short-lived trajectories and the (1-*f*) is the fraction of long-lived trajectories. We used *t* - 0.5 instead of *t* because the minimum trajectory duration is 0.5 s (10 frames) as previously mentioned. The residence time is the fitted slow lifetime (long-lived component). The dissociation rate constant (k_off_) used in the simulation is extracted from the confined residence time from VASH2-Halo.

### Correction for photobleaching

The photobleaching rate constant (*k*_photobleaching_) was calculated by fitting a single exponential decay function with an offset to the number of detected spots per frame over time.

The corrected slow dissociate rate constant (*k*_corrected_) is calculated as below:

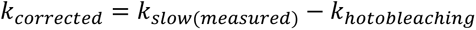

The residence time was corrected based on the corrected *k*_slow_ (slow dissociation rate constant). The statistics and fits for residence time and correction for photobleaching were performed using Graphpad Prism software.

### Computational Model

#### Monte Carlo model of Microtubule Dynamics

A Monte Carlo model was implemented to simulate an array of dynamic microtubules [57]. Microtubules grow and shrink by exchanging polymeric subunits with a soluble pool of tubulin, found in cells at a total concentration C_T_. All microtubules are nucleated from the boundary of a sphere of diameter d_center_ located in the center of a spherical shell of diameter d_cell_, representing the centrosome and outer boundary of our cell, respectively. The number of nucleation sites is set as N_T_, which also sets the maximum number of microtubules. At time t = 0, all microtubules are put in growth state, and they grow at a rate v_g0_ for a time Δt. After this initial step, each microtubule can either stay in growing state or a catastrophe event can happen, putting the microtubule in shortening state. Catastrophe events (growing to shortening transitions) happen to a growing microtubule with a frequency k_c_. For those microtubules that stay in growing state, the growth rate is considered as dependent on the free tubulin concentration after this initial step, v_g_ = k_on_C. As one micrometer of microtubule is made up of 1 µm·n_d_ dimers, C_T_ = C + 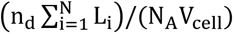, where L_i_ is the length of microtubule i, V_cell_ is the cell volume and N_A_ is the Avogadro number. microtubules keep growing until they either undergo a catastrophe or reach the cell boundary, where they immediately transition to shortening state. Shortening microtubules reduce their lengths at a rate v_s_ independent of tubulin concentration, and the shortening of microtubule i increases the free tubulin concentration by (n_d_ΔL_i_)/(N_A_V_cell_), where ΔL_i_ is the change in length of microtubule i. During its shortening state, a microtubule can either reach the centrosome of the cell or undergo a rescue event (shortening to growing transition). A microtubule which reaches the centrosome will completely depolymerize, and a new microtubule can be nucleated from this empty site with a frequency k_n_. Rescue events happen with a frequency k_r_ and they put a microtubule in growth state. All microtubules grow and shorten linearly, and their nucleation sites and directions of growth are selected at random during the first nucleation event. microtubules are modeled as cylinders of diameter d_t_, and their positions are selected to avoid intersections between microtubules or between microtubules and the centrosome. Two different sets of parameters for microtubule dynamics were employed in our simulations. Set B generates overly stable microtubules, which grow to long lengths and rarely depolymerize. Set A generates more dynamic microtubules, with lower tubulin concentrations, lower rescue frequencies and higher catastrophe frequencies, which better agree with experimental data [57]. Common parameters used to simulate microtubule arrays are shown in Table 2. Parameters associated to each of our microtubule sets are shown in Table 3. Schematics of our system and microtubule dynamic processes are shown in Figure 4. Comparison between the dynamics of sets A and set B are shown in Figure S8.

#### Reactive Brownian Dynamics

When a VASH enzyme is introduced in a cell containing an array of microtubules, the enzyme diffuses in the cell until it binds to one of the microtubules, which it can detyrosinate locally. VASH enzymes are modeled as spheres of diameter d_e_ which undergo 3D Brownian motion in the cell with a diffusion coefficient D_3D_. As enzymes cannot leave the cell, enter the centrosome, or traverse microtubules, special considerations must be taken to avoid these events. After a move is made at each time step Δt, the new location of the enzyme is checked to make sure that it follows these restrictions. If it does not, the location is rejected, and the enzyme is moved to a new location before the start of the next time step. When the enzyme either leaves the cell or enters the centrosome, this new location is determined by calculating the distance between the rejected location and the boundary in question and moving the enzyme along the vector going from the rejected location to the previous position with a magnitude equal to the measured distance. When the enzyme traverses a microtubule, it either binds to the tubule at the first intersection of the line defined by the previous and present positions and the microtubule with a probability p_bind_, or it instantly rebounds from the tubule with a probability 1 – p_bind_. In such case, the rebound is modeled akin to a rebound from the cell wall. A number N*_E_* of enzymes is randomly introduced in the system, and enzyme-enzyme interactions are neglected to keep the system computationally inexpensive.

When an enzyme successfully binds to a microtubule, it can slide along it in a diffusive (“Brownian”) manner, modeled by adding together a 1D Brownian motion with diffusion coefficient D_1D_ along the axis of the microtubule and a 1D rotational Brownian motion with diffusion coefficient D_θ_ along this same axis. Otherwise, it can stay confined in a certain tubulin subunit of length δ, modeled by a pure 1D rotational Brownian motion with diffusion coefficient D_θ_ along the axis of the microtubule. Directed modes of motion are neglected in this model. A molecule that binds to a microtubule can be found in Brownian (diffusive) mode with probability p_brown_, or in confined mode with probability p_conf_ = 1 – p_brown_. As a Brownian enzyme rarely becomes confined and vice versa, we will assume enzymes do not change modes during their bound state. The motion of Brownian enzymes can lead to escapes from the cell or into the centrosome. Hence, previously described boundary considerations are also applicable in this case. Brownian enzymes rebound at the end of the microtubules as well.

Another relevant process that enzymes may undergo after binding to a microtubule is unbinding, which can happen either randomly or as the result of the depolymerization of the subunit the enzyme is currently positioned in. The probability of unbinding differs based on the state of the enzyme. If the enzyme was diffusing along the microtubule, the probability of unbinding is given by p_unbrown_. On the other hand, if the enzyme was confined, the probability of unbinding is given by p_unconf_. After depolymerization, any enzyme located on a depolymerized subunit unbinds immediately. When an enzyme unbinds, it returns to its 3D Brownian motion regime, where it is again able to freely diffuse in all directions throughout the cell. The ejection of the enzyme from the microtubule is done along random direction, selected to avoid traversing the microtubule, over the same distance the enzyme would move were it freely diffusing in 3D.

#### Modeling Detyrosination

To detyrosinate a microtubule subunit, an enzyme must stay bound to it for a time *t_det_* ∼ 1.35 s based on its previously measured catalytic rate K_cat_ [33]. Molecules exhibiting confined motion stay bound to a subunit for an average time t_conf_ ∼ 3.03 s, while those exhibiting Brownian motion stay bound to a microtubule for an average time t_brown_ ∼ 1.56 s. During these ∼ 1.56 s, diffusing molecules move through many microtubule subunits very rapidly, staying on a particular one only for a 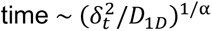, before moving to the next one, with *δ_t_* the length of the α-tubulin subunit (∼ 4 nm) and D_1D_ the coefficient of Brownian motion (MSD(t) = D_1D_t^α^). This time is much smaller than t_det_. Hence, we will assume only confined enzymes are capable of detyrosinating the subunit they are bound to, what they do with a probability 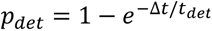. Note that, as *t_conf_* > *t_det_*, most confined enzymes are effectively capable of detyrosinating the subunit they are located at.

Once a subunit is detyrosinated, the dynamic properties associated to that subunit are modified, impacting the overall trends observed. This is modeled by changing the local frequencies and rates of a certain subunit *k_r_*, *k_c_*, *ν_g_*, *ν_s_* into *k_r_det__*, *k_c_det__*, *ν_g_det__*, *ν_s_det__*. The most significant changes involve the microtubule shortening properties. When a subunit is detyrosinated, catastrophe frequency and shortening rates decrease while growth parameters remain relatively invariant. As catastrophe events occur much less frequently, and shortening rates are significantly reduced, detyrosination leads to increased microtubule stability (see **Figure S8**).

Due to the polymeric nature of the microtubules, microtubule dynamics are dictated by the state of the subunit at the end of the microtubule, meaning that when a microtubule is in shortening state, its dynamics may change from subunit to subunit as they are removed (depolymerize). Subunits that have been detyrosinated cannot return to a tyrosinated state while they are polymerized in a microtubule [34]. The only way in which these subunits may return to the tyrosinated state is if they are depolymerized. Newly polymerized subunits are exclusively tyrosinated, meaning that after regrowth the previously detyrosinated subunit will now be in a tyrosinated state. For simplicity, we will consider each cylindrical microtubule to be split into uniformly distributed subunits bound by equidistant planes parallel to the cylinder’s base. As each micrometer of a microtubule has 1 µm * n_d_ subunits, our subunits will have a height of δ = 1/n_d_, or approximately 0.6 nm. This length is much smaller than the size of an α-tubulin subunit. However, we assume the detyrosination level will be well modeled using this treatment, as the number of subunits per micrometer remains constant.

#### Mimicking Experimental Resolution

Measuring the detyrosination level of each microtubule in our simulations is straight-forward, as we can keep track of the plus-end position of each microtubule and the tyrosination state of its subunits. The ratio between total detyrosinated length of a microtubule and its full length provides its detyrosination level. Averaging this ratio among all microtubules provides the total detyrosination level of our system. To compare this value to experiments, we reduced the resolution of the simulation system to that of DNA-PAINT experiments. The location of each microtubule subunit (whether detyrosinated or not) was blurred by jittering, with the purpose of generating a certain number of localizations per subunit, similarly to the ones from DNA-PAINT experiments. The number of localizations for each subunit was selected based on the log-normal probability that described the distribution of the number of localizations from single detyrosinated subunit from our experiments (**Figure 2E**), with μ= 2.7952 and standard deviation σ=0.64166. The directions of the jittered points are random and the distance of the jittered points from the true subunit position was randomly selected from a Gaussian distribution with full width at half maximum (FWHM) of 30 nm, which approximates our DNA-PAINT imaging resolution. For a system of sparse microtubules, where subunits associated with two different microtubules are unlikely to coexist in a single punctum, each punctum may include up to ∼30/0.6 = 50 subunits.

A detyrosinated punctum will then be equivalent to ∼50 detyrosinated sites, increasing the detyrosination level of a microtubule array. The detyrosination puncta and the total microtubule areas were defined by bounding points generated during resolution blurring described above. The detyrosination percentage was calculated by dividing the puncta area by the total microtubule area. Overall, at (relatively) low detyrosination levels (< 20% after blurring), the detyrosination level of our blurred microtubule arrays was well related to the “true” detyrosination level by multiplying it by a constant *χ* ∼ 50, as shown in **Figure S9 and S12**.

The previously described methodology applies both to **Model 1** and **Model 2**. For **Model 2**, when 20% of the maximum microtubule length is detyrosinated (percentage calculated after blurring to microscope resolution), the full microtubule gains the (more stable) kinetic properties of a detyrosinated microtubule over its full polymerized length.

## Acknowledgments

We thank Dr. Elena Sorokina, Dr. Melina Gyparaki, and Dr. Alexey Bogush for technical assistance, and we are grateful for reagents provided by Dr. Marie-Jo Moutin (University Grenoble Alpes) and Dr. Benjamin Prosser (University of Pennsylvania). We also thank Dr. Ekaterina Grishchuk, Dr. Michael Ostap, Dr. Benjamin Prosser, and Dr. Dimitrios Vavlylonis for critically reading the manuscript. This work is funded by R01 GM133842 (to M.L.), RM1 GM136511 (to M.L.) and CMMI-1548571 (to M.L.). Computational simulations used services provided by the OSG Consortium [81, 82], supported by the National Science Foundation awards #2030508 and #1836650, and by the Texas Advanced Computing Center (TACC) at The University of Texas at Austin.

