## Supporting Information for "Detyrosination enrichment on microtubule subsets is established by the interplay between a stochastically-acting enzyme and microtubule stability"

**Supplementary Figures**

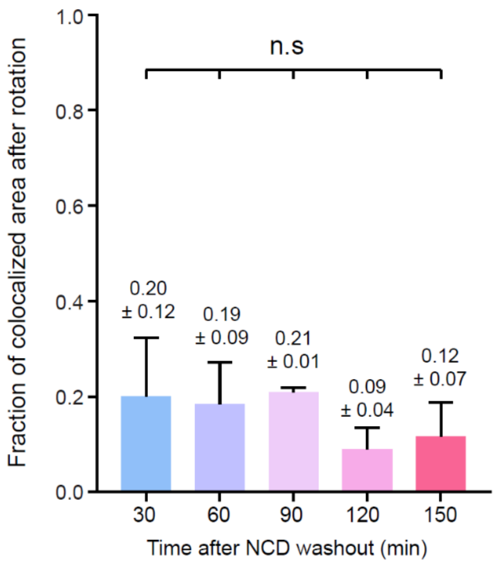


**Figure S1: Fraction of false positive puncta co-localized with microtubules.**

Each of the detyrosinated α-tubulin super resolution image was first flipped horizontally and again vertically with respect to the total β-tubulin image. The flipped image was then overlaid with the total β-tubulin image (original orientation) (Methods). The fraction of false positive co-localization is determined as the sum of the area of the flipped detyrosinated α-tubulin image co-localized with the β-tubulin image divided by the sum of the area of the original detyrosinated α-tubulin image co-localized with the β-tubulin image. The mean and SD of independent experiments are shown for each time point after Nocodazole washout. For each time point after Nocodazole washout, 30 min, $N$=15 cells from 5 experiments; 60 min, $N$ =17 cells from 4 experiments; 90 min, $N$ = 17 cells from 3 experiments; 120 min, $N$ = 13 cells from 3 experiments; 150 min, $N$ = 9 cells from 3 experiments. One-way ANOVA and Tukey's multiple comparison was performed. $P$ >0.05, n.s, not significant.

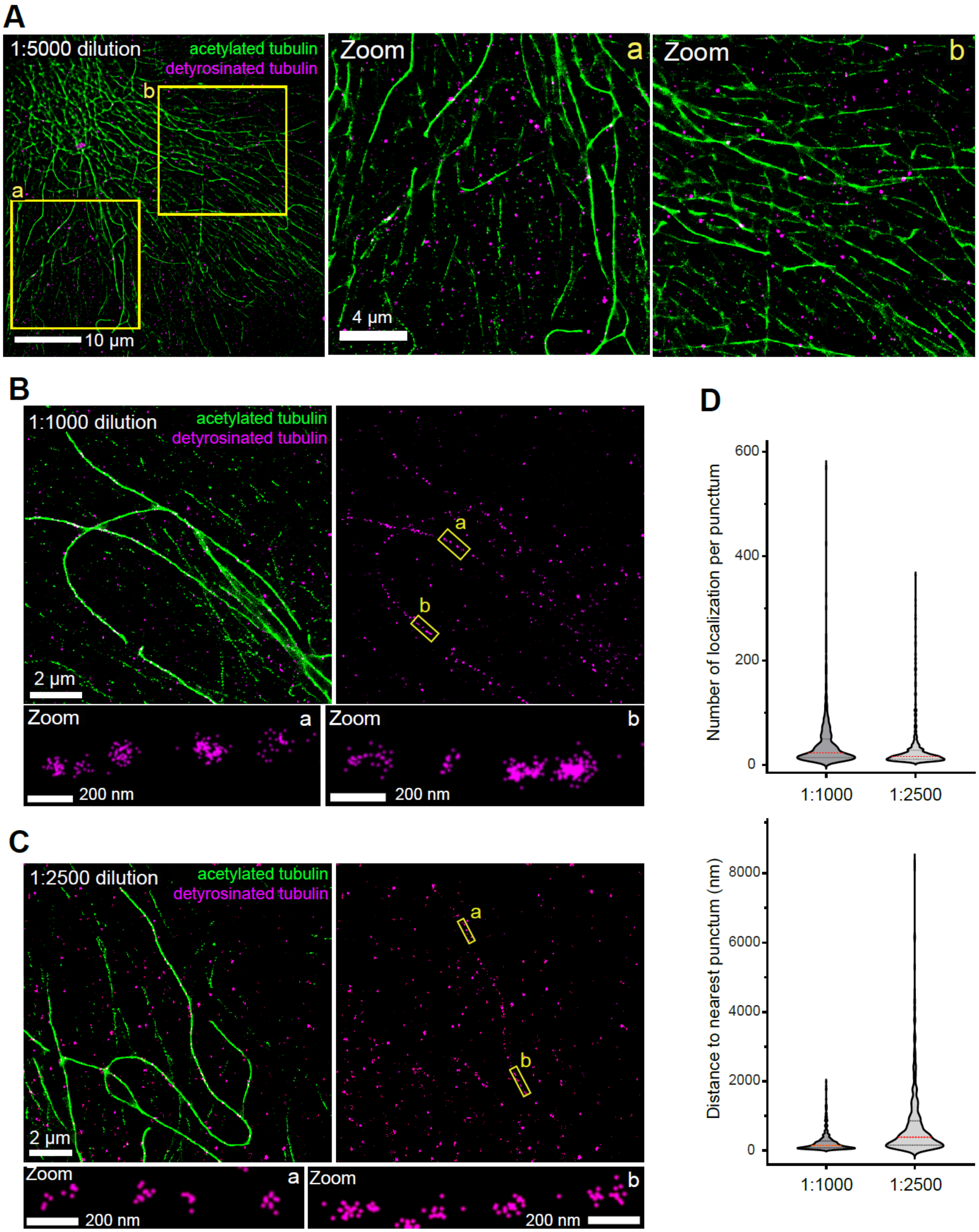


**Figure S2: Comparison of different dilutions of anti-detyrosinated α-tubulin antibody to detect single detyrosination sites.**

(**A-C**) Representative two-color DNA PAINT image showing acetylated microtubules (green) and detyrosinated microtubules (magenta) stained with anti detyrosinated α-tubulin antibodies at 1:5000 dilution (**A**), 1:1000 dilution (**B**) and 1:2500 dilution (**C**). Two regions (a, b, yellow boxes) for each example are selected for zoom. Scale bars in (**A**), 10 µm; (**B-C**), 2 µm. Scale bars in zoom, 4 µm (**A**), 200 nm (**B-C**). Detyrosination on microtubules are undetectable at 1:5000 dilution of the primary antibodies, and the most prominent signal appears as non-specific background puncta, which do not overlap with microtubules. (**D**) Violin plots showing the comparisons of the number of localizations per puncta (upper plot) and distance to nearest puncta (lower plot) detected by 1:1000 or 1:2500 dilution of the anti-detyrosinated α-tubulin antibodies. The red dash lines indicate the median value, the upper black dash lines are the 75$^{th}$ percentile, the lower black dash lines show the 25$^{th}$ percentile. The 1:2500 dilution of the primary antibodies allows for detection of primarily single detyrosination sites as shown by the larger nearest neighbor distance (median: 386 nm), the narrower distribution and lower number of localizations per puncta (median: 16) as compared to the 1:1000 dilution (median: 147 nm nearest punctum distance and median 23 localizations per punctum).

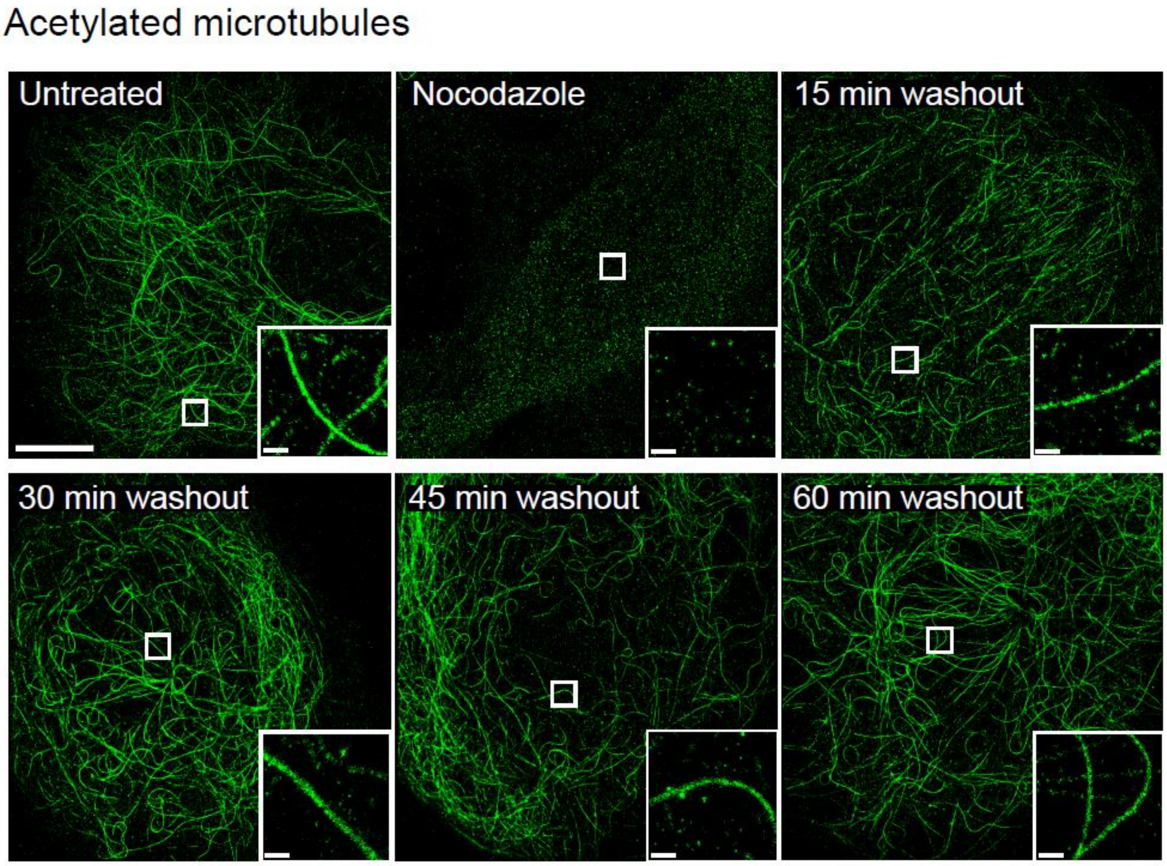


**Figure S3: Acetylation is established rapidly after Nocodazole washout.**

Representative DNA-PAINT super resolution images showing acetylated microtubules in BSC-1 cells that are untreated, treated with 33 µM Nocodazole for 3 hr at 37 ºC, and at 15, 30, 45, 60 min after Nocodazole washout. Acetylation was already re-established at 30-60-min post washout. Scale bar, 10 µm. Insets are the zoomed regions (open squares). Scale bars in zooms, 500 nm.

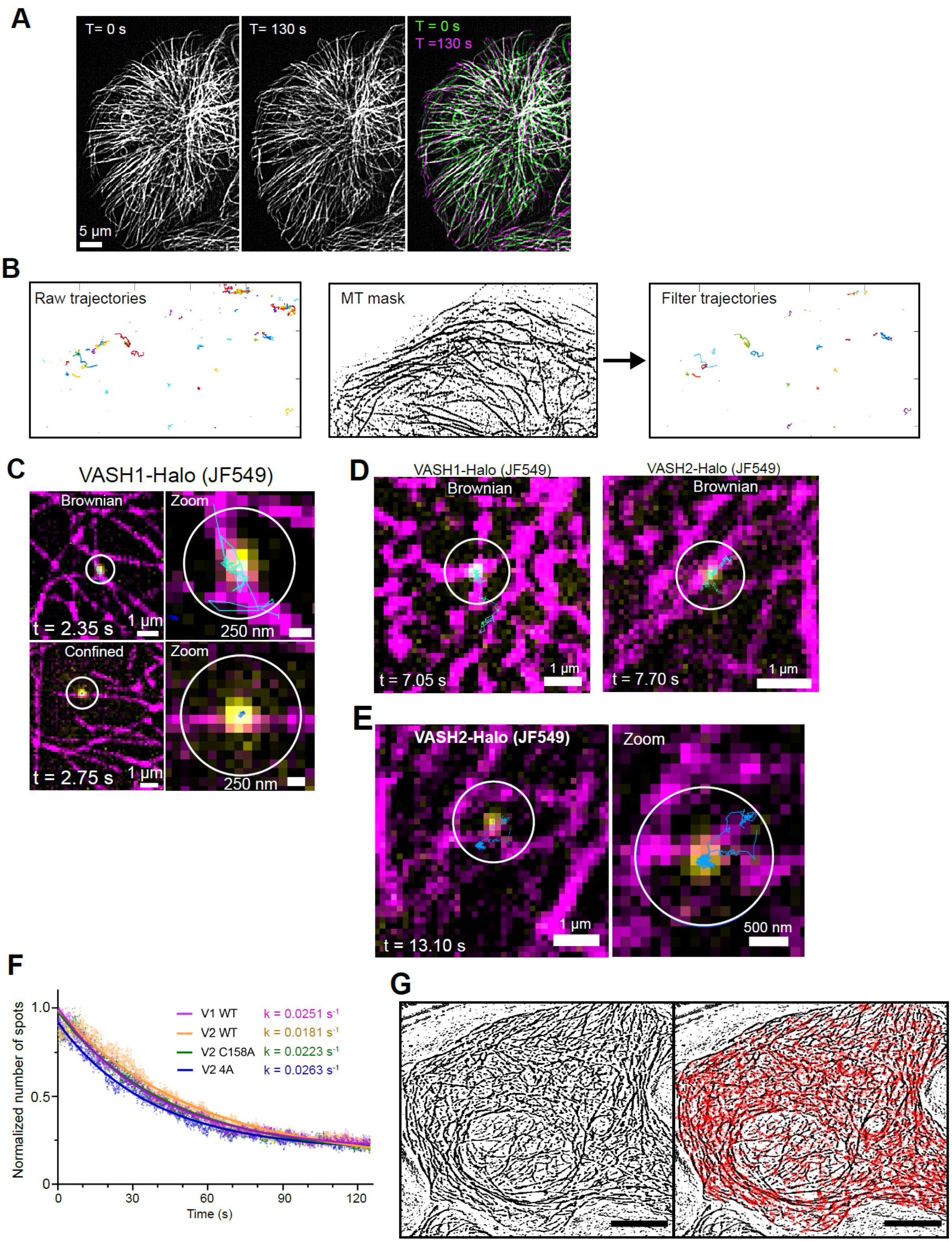


**Figure S4: Example motion trajectories of VASH1- and VASH2-Halo+JF549 on microtubules.**

(**A**) Representative confocal images of microtubule network in U2OS cells labeled with Tubulin Tracker Deep red at 0 s and 130 s. The majority of the microtubules are overlapping but modest fluctuations occurred (merge). (**B**) Example showing how trajectories were filtered based on co-localization with the microtubule mask generated by binarization of confocal snapshots of microtubules labeled by tubulin tracker. **(C)** Representative overlay images showing trajectories of single VASH1-Halo molecules labeled with JF549 ligand (yellow) on confocal snapshot images of microtubules labeled with Tubulin Tracker (magenta) in live U2OS cells. Scale bars, 1 µm. Scale bars in zoom, 250 nm. (**D**) Examples of prolonged diffusion of VASH1- and VASH2-Halo at microtubule-dense area where microtubules crisscross. The duration (t) of the trajectories are shown. Scale bars, 1 µm. (**E**) An example trajectory showing a VASH2-Halo molecule transition between Brownian and confined motions. Scale bar, 1 µm. Scale bar in zoom, 500 nm. (**F**) The photobleaching rate constants were calculated based on the number of spots detected at each frame over time. The number of spots per frame was normalized as a fraction to the highest number of spots among all the frames. A single exponential decay was fitted to each data set to obtain the decay rate constants. (**G**) An example of VASH2-Halo trajectories overlapped with binarized images of microtubules in a cell. Scar bars, 10 µm.

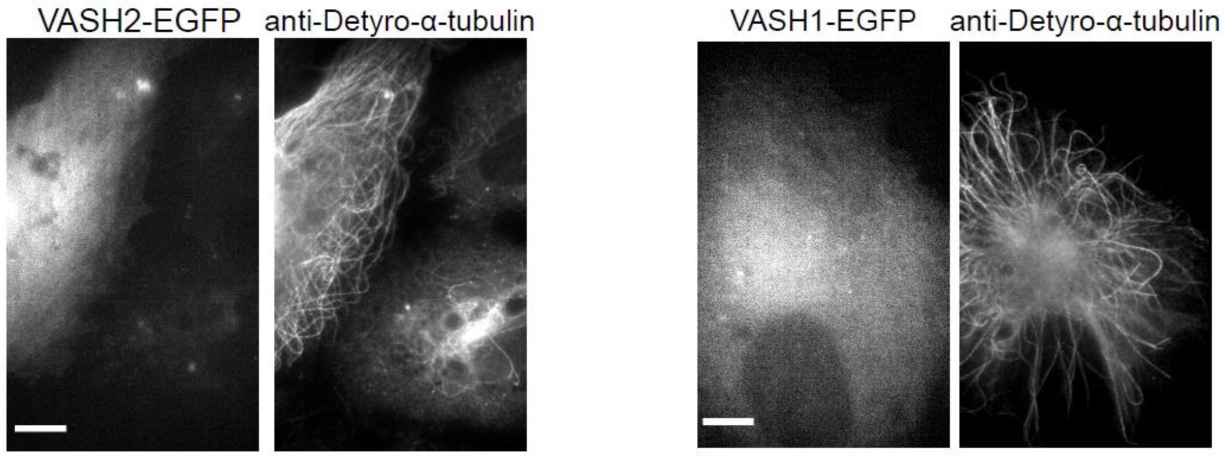


**Figure S5: VASH1 and VASH2 are cytosolic proteins.**

Wide field epi-fluorescent images showing ectopic expression of VASH1- or VASH2-EGFP (each co-expressed with SVBP-Myc-FLAG) in BSC-1 cells. The EGFP (left) signal is mostly diffuse in the cytosol. Scale bars, 10 µm.

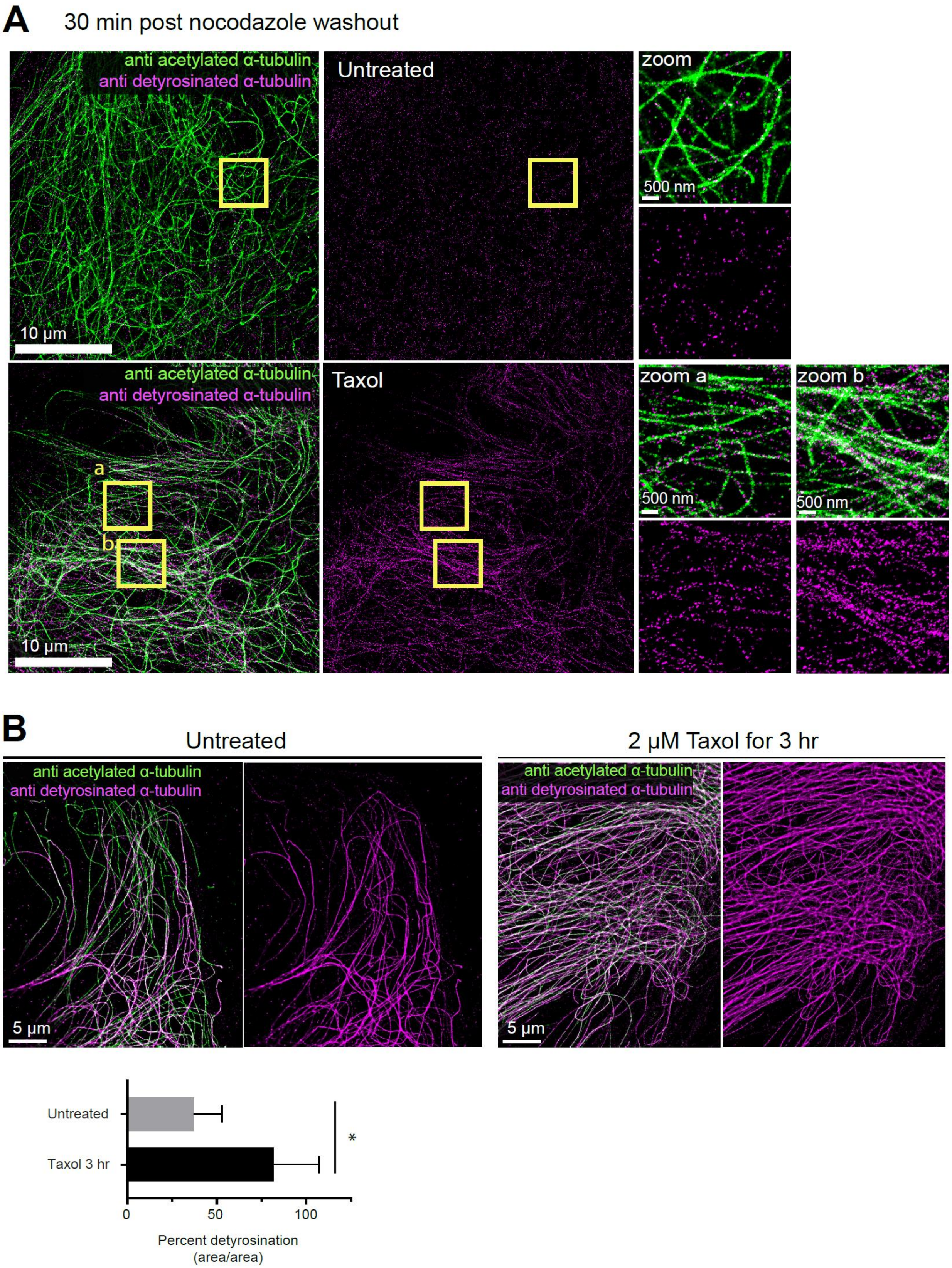
A
30 min post nocodazole washout
anti acetylated α-tubulin
anti detyrosinated α-tubulin
Untreated
zoom
500 nm
10 μm
Taxol
zoom a
zoom b
a
b
B
Untreated
2 μM Taxol for 3 hr
5 μm
Untreated
Taxol 3 hr
*
0
50
100
Percent detyrosination
(area/area)

**Figure S6: Detyrosination increases significantly upon Taxol stabilization of microtubules.** (**A**) Representative two-color DNA PAINT images showing nascent detyrosination in BSC-1 cells that are untreated (DMSO) or treated with 2 µM Taxol for 30 min at 37ºC, immediately after the removal of 33 µM Nocodazole treatment for 3 hr. Detyrosination puncta (magenta) appear much denser on all microtubules (green) upon Taxol treatment. Regions selected for zoom are indicated by yellow boxes. Scale bars, 10 µm. Scale bars in zoom, 500 nm. (**B**) Representative two-color DNA-PAINT images showing detyrosination in BSC-1 cells incubated with 0.04% DMSO (untreated) or 2 µM Taxol for 3 hr at 37ºC. Detyrosination appears prevalent on most microtubules upon Taxol treatment. Scale bars, 5 µm. Detyrosination percentage calculated based on area, increased from 37% ± 15% (untreated) to 82% ± 25% (Taxol 3 hr). Unpaired t-test with Welch's correction was performed. * $p < 0.05$.

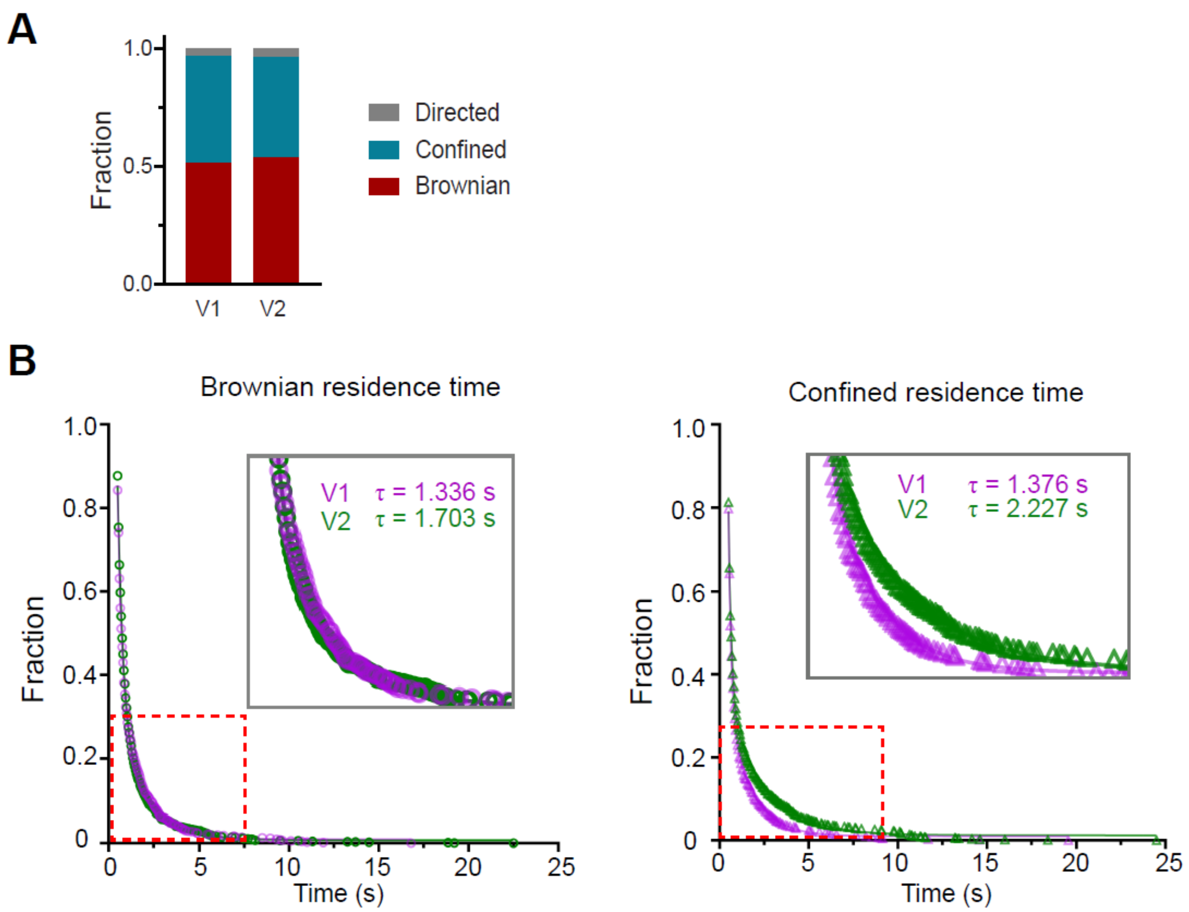


**Figure S7: Single molecule tracking of VASH at 30 min post Nocodazole washout**

Single molecule tracking of VASH1-Halo (V1) and VASH2-Halo (V2) at 30 min post Nocodazole washout when microtubules are mostly tyrosinated. (**A**) Fraction of different classes of motion for VASH1 (V1) and VASH2 (V2). (**B**) Double exponential fits for calculating residence time for V1 (magenta) and V2 (green). The residence times ($\tau$) for V1 and V2 in each motion class are shown. The 95% confidence intervals are shown in Table 1.

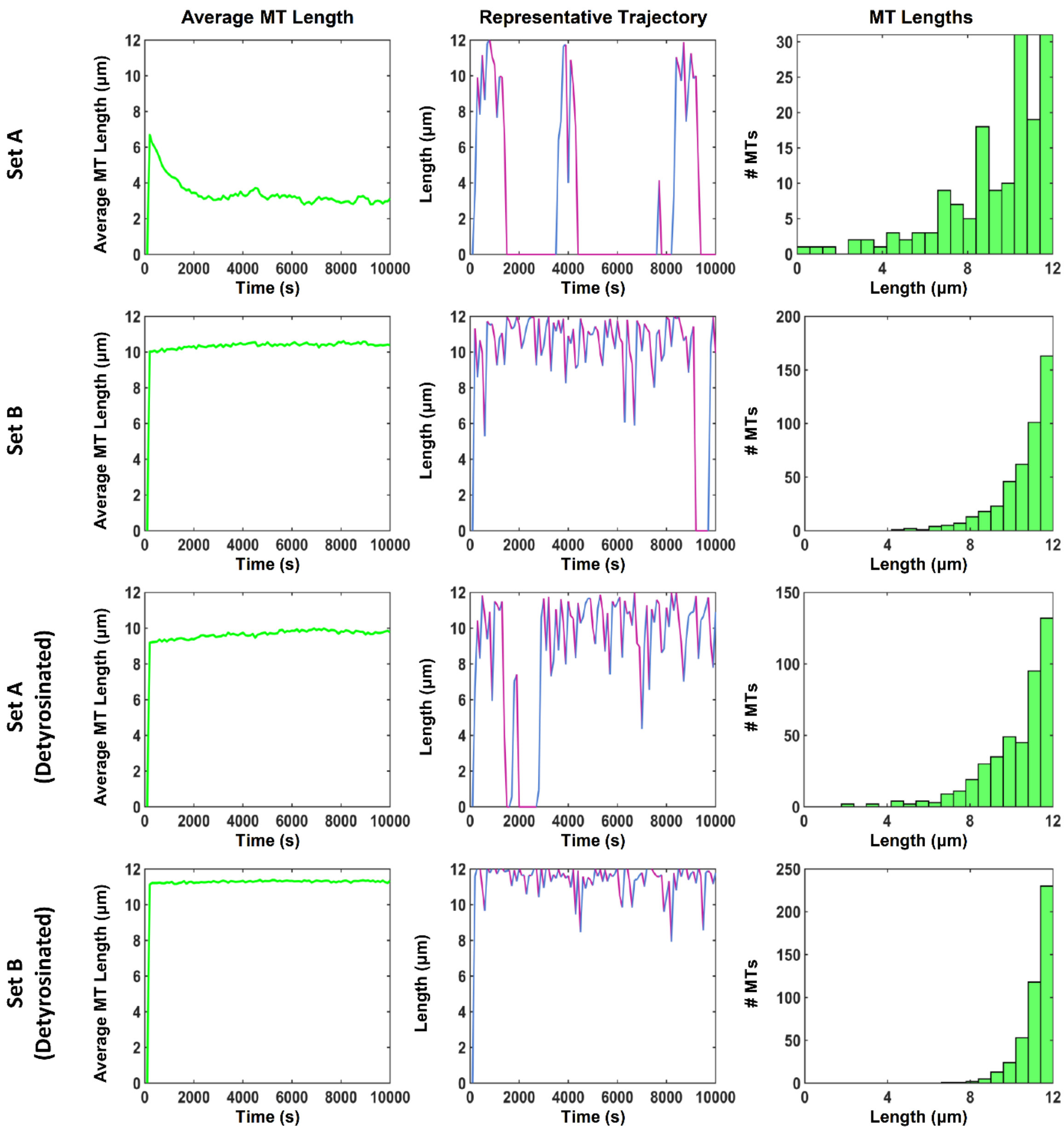


**Figure S8: Comparisons between microtubule dynamics for highly dynamic and intermediately dynamic arrays in the absence of enzymes ($N_T = 500$). (LEFT)** Average microtubule lengths evaluated after 10,000 s for tyrosinated microtubules (two top rows) and for microtubules with parameters modified to represent permanently detyrosinated microtubules ($k_{r_{det}}$, $k_{c_{det}}$, $v_{g_{det}}$ and $v_{s_{det}}$) (two bottom rows). Average lengths: $L_T = 3.39 \pm 4.69$, $L_T = 10.27 \pm 2.55$, $L_T = 9.56 \pm 3.29$ and $L_T = 11.18 \pm 1.47$ for highly dynamic, intermediately dynamic, highly dynamic but detyrosinated, and intermediately dynamic but detyrosinated microtubule arrays.

**(MIDDLE)** Example length trace over time of a single microtubule. **(RIGHT)** Distribution of microtubule lengths evaluated after 10,000 s.

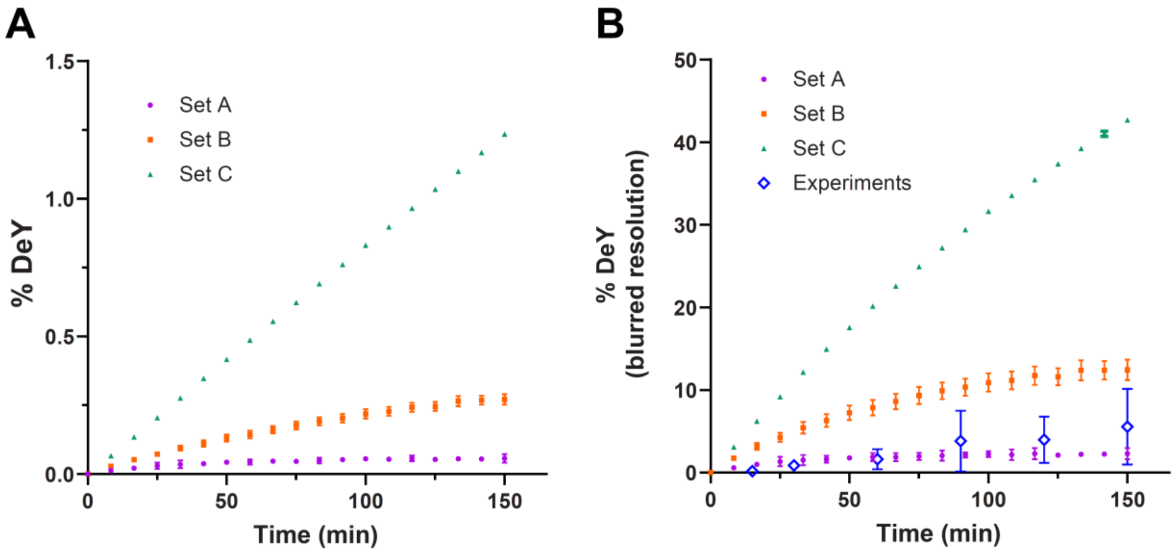


**Figure S9: Early-stage detyrosination accumulation kinetics simulated using Model 1 simulations. (A)** Simulated detyrosination levels (symbols) as a function of time for 250 microtubules that are highly dynamic (set A), intermediately dynamic (set B) and fully stable (set C) with 100 nM VASH. **(B)** Comparison between experimental detyrosination levels (open blue diamonds, ± SD) and simulated detyrosination levels presented in (A) after blurring to experimental resolution. Set A microtubules most closely mimicked the detyrosination levels we observed in the DNA-PAINT experiment. All simulated values are shown as mean ± SD.

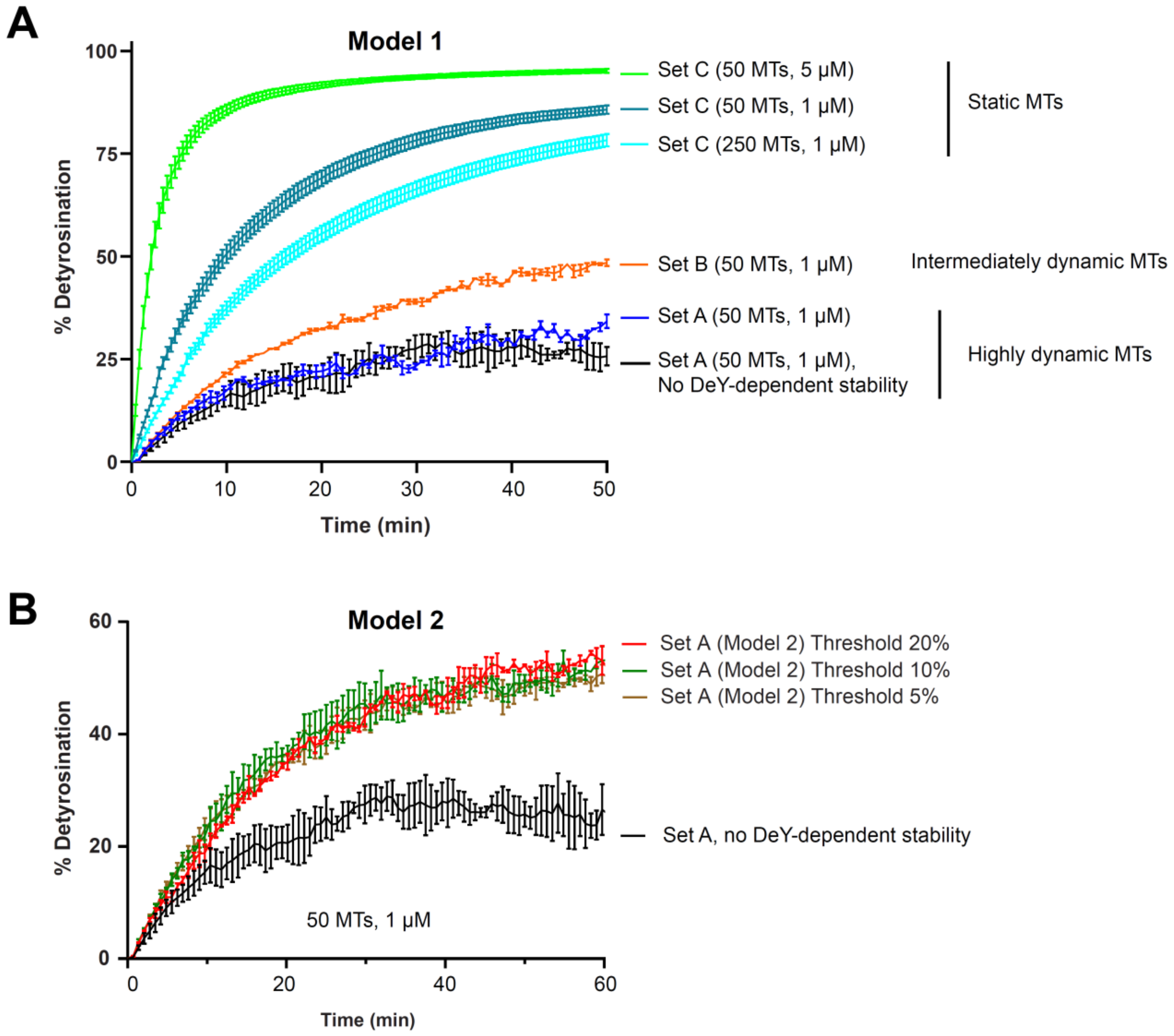


**Supplementary Figure 10: Steady state detyrosination. (A)** Under Model 1**,** steady states were reached by simulating 1-5 μM VASH with 50 or 250 microtubules that are highly dynamic (set A), intermediately dynamic (set B), or fully stable (set C). Detyrosination levels saturated at ~ 23% (purple) and ~ 52% (orange) for highly and intermediately dynamic microtubules respectively. Stable microtubules reached detyrosination levels of ~ 94% (navy blue), ~ 97% (light green) and ~ 88% (cyan), which fall outside the margins of the plot. The black curve corresponds to the control Model in which detyrosination does not change microtubule stability. A higher concentration of VASH enzymes was used to examine steady state levels of detyrosination under reasonable simulation times. Note that, as expected from our experiments using Taxol to stabilize microtubules (**Figure S6**), we observe that higher initial microtubule stability leads to faster detyrosination as well as higher plateau levels of detyrosination at steady state. Also, note the scalings with the number of microtubules and of enzymes: higher numbers of enzymes lead to higher detyrosination levels, while the opposite is observed for the numbers of microtubules. **(B)** Under Model 2 (threshold-based), steady state detyrosination levels in highly dynamic microtubules became significantly higher (~52 %) compared to no detyrosination-dependent stability control (~23 %). Note that the same no detyrosination-dependent stability control simulation was included in both **(A)** and **(B)** for comparison. Error bars represent mean ± SD of three simulations.

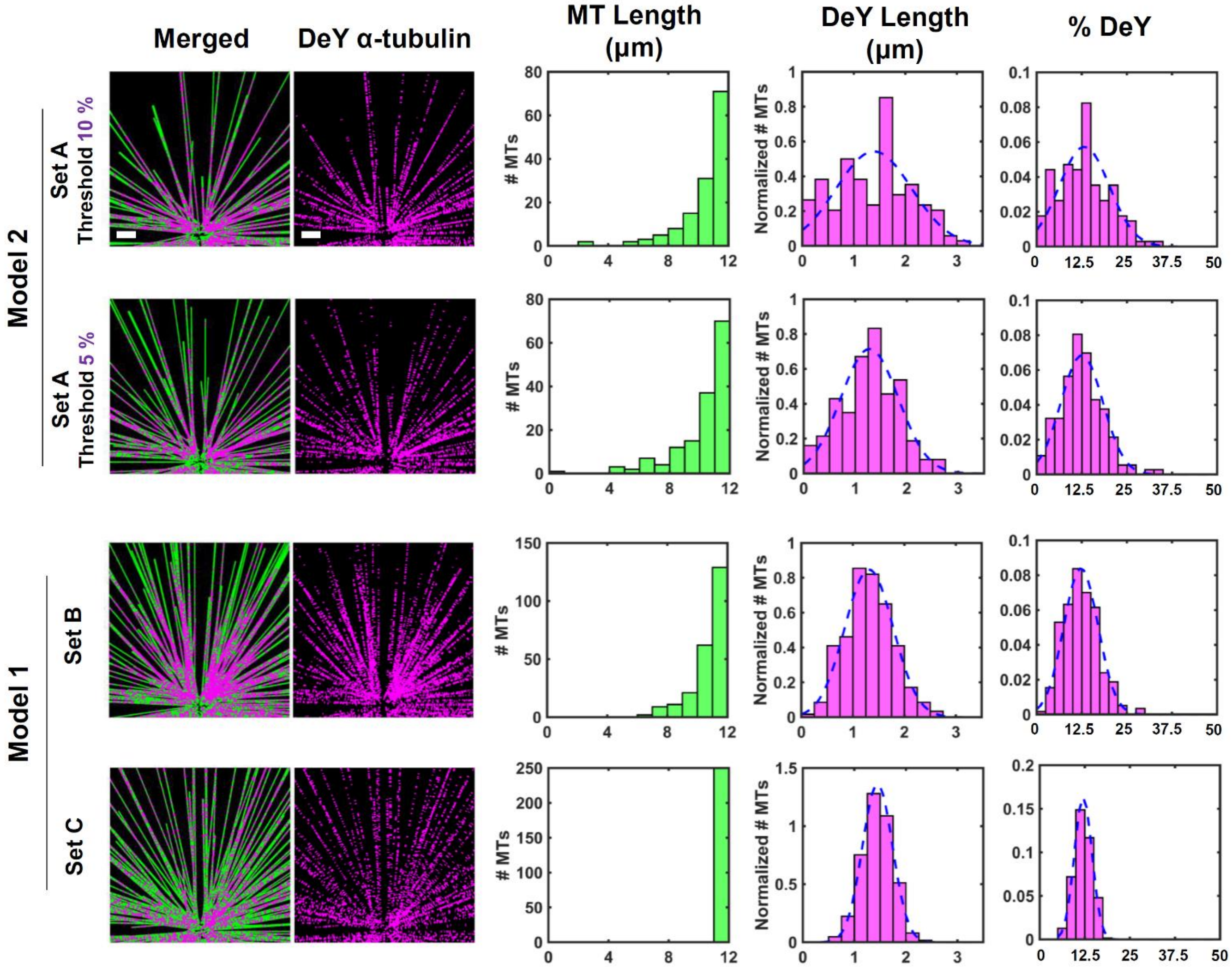


**Supplementary Figure 11: Detyrosination distributions under different stability models. (LEFT)** Representative images showing regions of microtubules at 12% total detyrosination with 500 nM VASH. Detyrosinated α-tubulin subunits are shown in magenta and β-tubulin in green. The thresholds of detyrosination percentages (in purple letters) for microtubule stability transition for Model 2 are indicated. Scale bars, 2 µm. Set A, Set B and Set C denote the inherent microtubule dynamic regimes as highly dynamic, intermediately dynamic and fully stable (**Table 3**). **(RIGHT)** Histograms of the total microtubule length, detyrosinated segment length, and percentage of detyrosination. Fully depolymerized microtubules were neglected in our histograms. Percentages of detyrosination were best fitted by single Gaussian distribution (blue dashed lines). The mean (µ) and variance (σ) from the gaussian fits are reported in **Supplementary Table 1.**

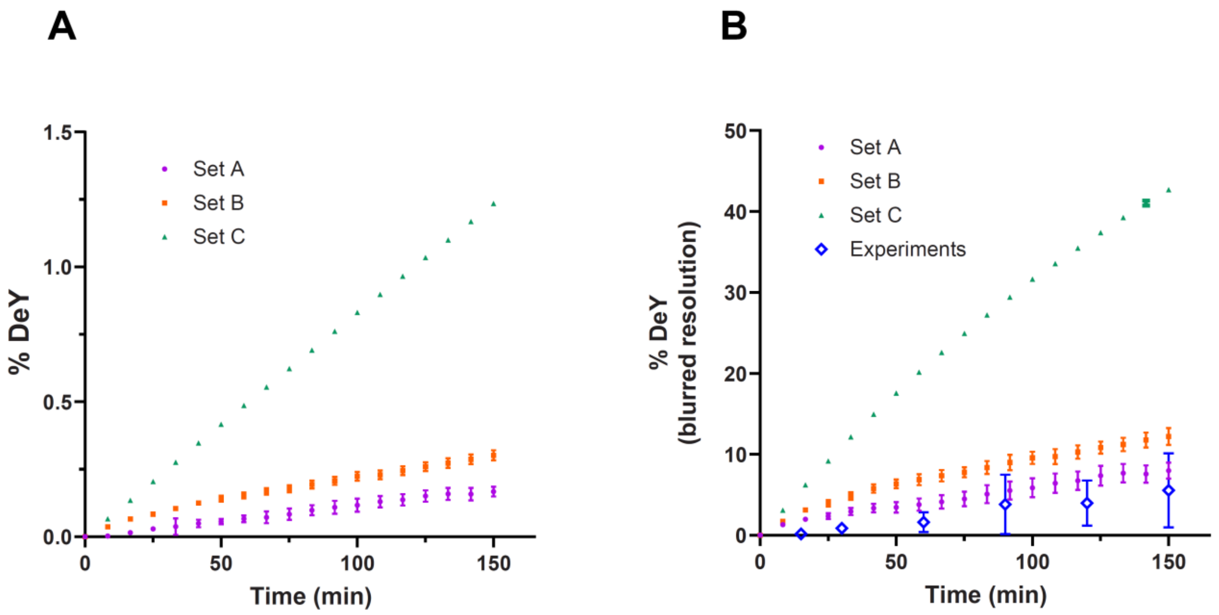


**Figure S12: Early-stage detyrosination accumulation kinetics simulated using Model 2 (20% threshold). (A)** Simulated detyrosination levels (symbols) as a function of time for 250 microtubules that are highly dynamic (set A), intermediately dynamic (set B) and fully stable (set C) with 100 nM VASH. **(B)** Comparison between experimental detyrosination levels (open dark blue diamonds, ± SD) and simulated detyrosination levels presented in (A) after blurring to experimental resolution. All simulated values are shown as mean ± SD. Set A microtubules most closely mimicked the detyrosination levels we observed in the DNA-PANT experiments.

**Supplemental Tables**

**Supplemental Table S1.** Mean (μ), variance (σ) and amplitude (C) of gaussian fits on detyrosination distribution on highly dynamic microtubules (set A) under different detyrosination-dependent stability models.

| Dynamics and model | Best fit | | DeY length (μm) | % DeY |
|---|---|---|---|---|
| Set A, no DeY-dependent stability | double Gaussian | $\mu_1$ | 1.4729 | 22.54 |
| | | $\mu_2$ | 0.4982 | 9.1852 |
| | | $\sigma_1$ | 0.388 | 123.6378 |
| | | $\sigma_2$ | 0.078 | 29.5869 |
| | | $C_1$ | 0.5614 | 0.3171 |
| | | $C_2$ | 0.4386 | 0.6829 |
| Set A (**Model 1**) | double Gaussian | $\mu_1$ | 0.6191 | 8.6538 |
| | | $\mu_2$ | 1.5292 | 22.0317 |
| | | $\sigma_1$ | 0.1116 | 24.2568 |
| | | $\sigma_2$ | 0.4733 | 62.0527 |
| | | $C_1$ | 0.4973 | 0.6627 |
| | | $C_2$ | 0.5027 | 0.3373 |
| Set A (**Model 2**, 20 % threshold) | double Gaussian | $\mu_1$ | 0.8683 | 8.0868 |
| | | $\mu_2$ | 2.7318 | 22.6361 |
| | | $\sigma_1$ | 0.251 | 19.8803 |
| | | $\sigma_2$ | 0.092 | 30.1382 |
| | | $C_1$ | 0.8697 | 0.6897 |
| | | $C_2$ | 0.1303 | 0.3103 |
| Set A (**Model 2**, 10 % threshold) | Gaussian | $\mu$ | 1.3665 | 12.9966 |
| | | $\sigma$ | 0.7266 | 6.9105 |
| Set A (**Model 2**, 5 % threshold) | Gaussian | $\mu$ | 1.2807 | 12.5284 |
| | | $\sigma$ | 0.5515 | 5.688 |
| Set B (**Model 1**) | Gaussian | $\mu$ | 1.3132 | 12.2764 |
| | | $\sigma$ | 0.4726 | 4.774 |
| Set C (**Model 1**) | Gaussian | $\mu$ | 1.4502 | 12.0601 |
| | | $\sigma$ | 0.2934 | 2.4401 |
